# Mitotic retention of H3K27 acetylation promotes rapid topological and transcriptional resetting of stem cell-related genes and enhancers upon G1 entry

**DOI:** 10.1101/2020.06.02.130104

**Authors:** Bobbie Pelham-Webb, Alexander Polyzos, Luke Wojenski, Andreas Kloetgen, Jiexi Li, Dafne Campigli Di Giammartino, Leighton Core, Aristotelis Tsirigos, Effie Apostolou

## Abstract

The identity of dividing cells is challenged during mitosis, as transcription is halted and chromatin architecture drastically altered. How cell type-specific gene expression and genomic organization are faithfully reset upon G1 entry in daughter cells remains elusive. To address this issue, we characterized at a genome-wide scale the dynamic transcriptional and architectural resetting of mouse pluripotent stem cells (PSCs) upon mitotic exit. This revealed distinct patterns of transcriptional reactivation with rapid induction of stem cell genes and their enhancers, a more gradual recovery of metabolic and cell cycle genes, and a weak and transient activation of lineage-specific genes only during G1. Topological reorganization also occurred in an asynchronous manner and associated with the levels and kinetics of transcriptional reactivation. Chromatin interactions around active promoters and enhancers, and particularly super enhancers, reformed at a faster rate than CTCF/Cohesin-bound structural loops. Interestingly, regions with mitotic retention of the active histone mark H3K27ac and/or specific DNA binding factors showed faster transcriptional and architectural resetting, and chemical inhibition of H3K27 acetylation specifically during mitosis abrogated rapid reactivation of H3K27ac-bookmarked genes. Finally, we observed a contact between the promoter of an endoderm master regulator, *Gata6*, and a novel enhancer which was preestablished in PSCs and preserved during mitosis. Our study provides an integrative map of the topological and transcriptional changes that lead to the resetting of pluripotent stem cell identity during mitotic exit, and reveals distinct patterns and features that balance the dual requirements for self-renewal and differentiation.

---

Mitosis (MIT) poses a temporal challenge for cell identity, since it is accompanied by global transcriptional silencing (Gottesfeld and Forbes, 1997; Taylor, 1960), dissociation of most transcription factors and cofactors from target genes (Martínez-Balbás et al., 1995) and profound reorganization of 3D chromatin architecture (Earnshaw and Laemmli, 1983; Marsden and Laemmli, 1979). Under self-renewing conditions, faithful propagation of cell identity relies on the proper reestablishment of cell-type defining molecular features during mitotic exit and entry to the next Gap 1 (G1) phase. Therefore, MIT-to-G1 transition is a period of critical importance for cell fate decisions (Boward et al., 2016; Soufi and Dalton, 2016) and a unique time window to study the principles of transcriptional and architectural resetting.

Previous work in human and mouse cell lines has started to dissect the 3D chromatin reorganization during the cell cycle and described distinct models of mitotic or interphase chromosome folding (Gibcus et al., 2018; Liang et al., 2015b; Naumova et al., 2013; Ou et al., 2017) as well as dynamic reorganization during mitotic exit (Abramo et al., 2019; Dileep et al., 2015; Nagano et al., 2017; Zhang et al., 2019). Unrelated studies have reported distinct waves of transcriptional reactivation during mitotic release and a global, transient spike in transcription during G1 entry (Hsiung et al., 2016; Palozola et al., 2017). However, the interplay between architectural resetting and transcriptional reactivation during mitotic exit, which likely holds the key how cell identity is maintained, remains unclear. Similarly, the mechanisms responsible for the differential kinetics of molecular resetting and the potential significance for regulation of cell identity are yet to be uncovered.

One proposed mechanism for rapid molecular resetting and faithful heritability of cell identity is mitotic bookmarking, which refers to the partial retention of selected factors and features on mitotic chromatin (Michelotti et al., 1997).Although several mitotically retained features have been described, including DNA and histone modifications (Margueron and Reinberg, 2010; Zaidi et al., 2010), epigenetic modulators (Blobel et al., 2009; Dey et al., 2009), and DNA-binding transcription factors (TFs) (Caravaca et al., 2013; Deluz et al., 2016; Festuccia et al., 2016; Kadauke et al., 2012; Liu et al., 2017b; Teves et al., 2018; Teves et al., 2016), only few have been directly tested and validated to have a functional role in the resetting and/or maintenance of cell identity. Recent work using advanced genomics and proteomics technologies has further challenged the extent of transcriptional and epigenetic shutdown during mitosis. Acetylation of several histone tail residues were shown to persist on mitotic chromatin both globally and at specific genomic sites (Behera et al., 2019; Javasky et al., 2018; Liu et al., 2017a; Liu et al., 2017b); chromatin accessibility was found to be largely unchanged in mitosis, especially at promoters (Hsiung et al., 2015; Teves et al., 2016), and transcription itself to continue at low levels (Liang et al., 2015a; Liu et al., 2017a; Palozola et al., 2017).

To what degree mitotic retention of one or more of these features promotes faster and faithful reestablishment of the cell type-specific transcriptional program and 3D chromatin landscape upon G1 entry remains largely unexplored.

MIT-to-G1 transition has been shown to be particularly critical for the regulation of pluripotent stem cell (PSC) identity (Soufi and Dalton, 2016). PSCs hold great promise for regenerative medicine and disease modeling, since they are characterized by the capacity to self-renew indefinitely in culture while maintaining their ability to differentiate into all somatic cell types upon proper stimulation (Evans, 2011; Tabar and Studer, 2014). The PSC cell cycle is very rapid (10-12 hours in mouse and 16 hours in humans) with an exceptionally short G1 phase, compared to most somatic cell types (Savatier et al., 2002), which is the critical window for PSCs to “decide” either to self-renew or respond to differentiation cues towards defined lineages (Coronado et al., 2013; Pauklin et al., 2016; Pauklin and Vallier, 2013; Sela et al., 2012; Singh et al., 2015). Therefore, studying the mechanisms and principles of molecular resetting during MIT-to-G1 in PSCs might aid the rationale optimization for biomedical applications using these cells, such as directed differentiation.

In this study, we used PRO-seq and *in situ* Hi-C in mouse PSCs to characterize for the first time the transcriptional and architectural landscape during mitosis and to capture the order of molecular events that result in the reestablishment of cell identity upon mitotic exit. Our integrative analysis allowed us to identify distinct patterns of transcriptional and topological resetting and provide insights into their temporal interconnections and their relevance for PSC identity. Moreover, we discuss potential mechanisms that dictate differential recovery rates and provide strong evidence for the role of mitotic bookmarking in promoting both faster transcriptional and architectural resetting upon mitotic exit. Together, these results provide insight into the molecular logic that underlies the reestablishment of cell type-specific gene expression and chromatin organization during mitotic exit in PSCs.

## RESULTS

### Mitotic arrest and release of pluripotent stem cells into G1

In order to characterize the molecular resetting of mouse pluripotent stem cells (PSCs) after cell division, we performed a mitotic release time course (Figures 1A and S1A). PSCs arrested in mitosis (MIT) were isolated after nocodazole treatment followed by mitotic shake-off, as previously described (Liu et al., 2017b). Mitotic purity was verified by staining with H3Ser10p (Hendzel et al., 1997) followed by Fluorescence Activated Cell Sorting (FACS) (Figures 1B) and immunofluorescence analysis (Figure 1C), and cell preparations with <95% purity were discarded. After extensive nocodazole wash off, mitotic cells were released in full medium and collected at different time intervals for evaluation of their DNA content (Figure 1D) and cell cycle fluorescent markers using the FUCCI2a system (Figure 1E) (Mort et al., 2014). Multiple independent experiments indicated that the majority of cells entered G1 around 1 hour after release (55%-70% 2N population) and that G1 ended 2-3 hours later when DNA replication began (Figures S1B and S1C), in line with previous studies in mouse PSCs (Coronado et al., 2013). Therefore, we chose the 1 hour and 3 hour time points to represent early G1 (EG1) and late G1 (LG1), respectively. Asynchronous (ASYN) PSCs, comprised mostly by cells in S/G2 phase (>70%) (Figure 1E) were analyzed to represent a later stage of resetting. For each time point (MIT, EG1, LG1 and ASYN), biological replicates from two independent experiments with similar purity and release kinetics were collected for Precision nuclear run-on sequencing (PRO-seq) (Kwak et al., 2013) to track the genome-wide resetting of nascent transcriptional activity and for *in situ* Hi-C (Rao et al., 2014) to capture the global architectural reorganization (Figure 1A).

**Figure 1.**
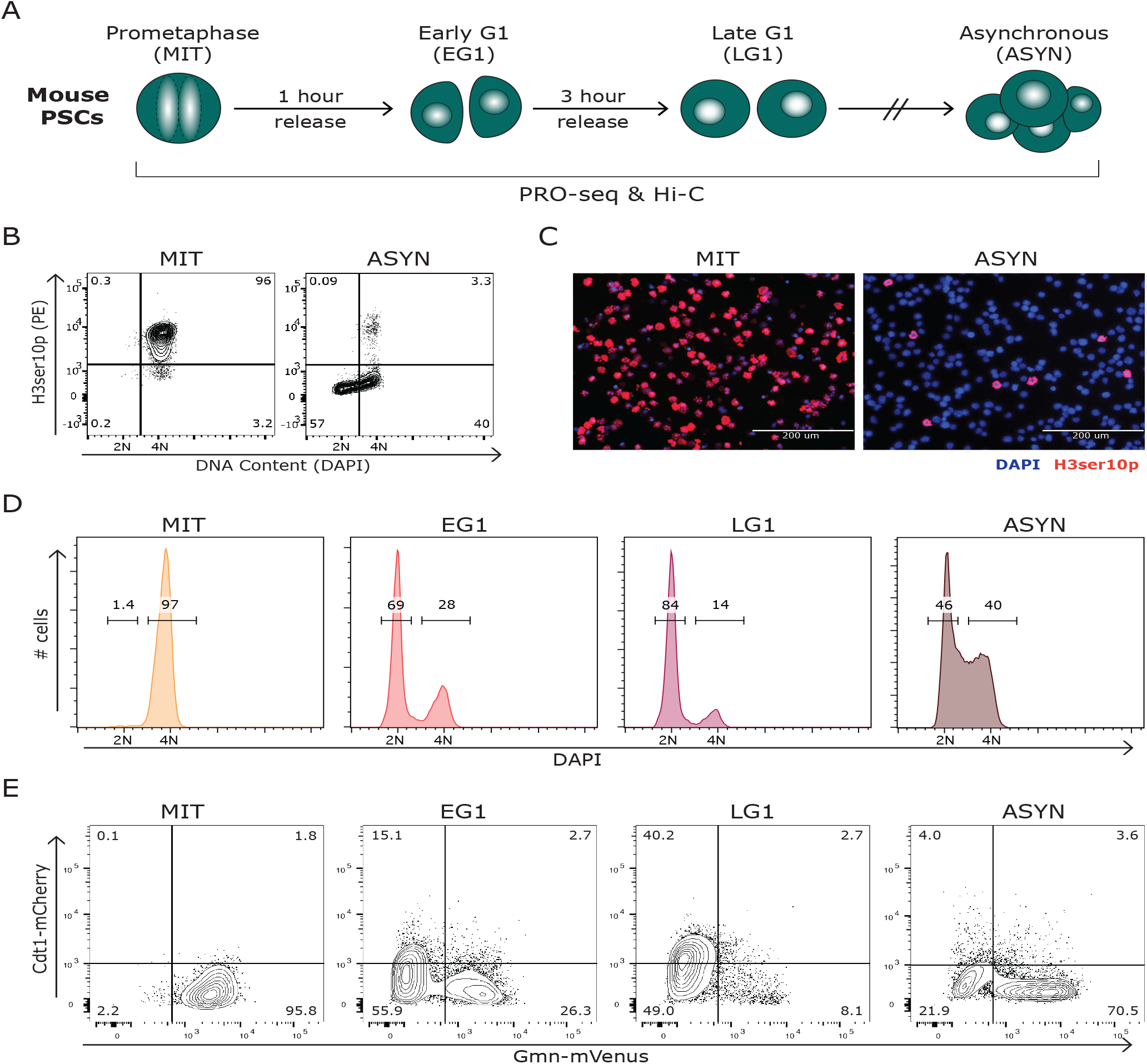
Mitotic arrest and release of pluripotent stem cells into G1. (A) Schematic depicting our strategy for collecting and profiling by PRO-seq and Hi-C mouse pluripotent stem cells (PSCs) in Mitosis (MIT), Early G1 (EG1), Late G1 (LG1), and untreated, asynchronous cells (ASYN). (B) FACS plots showing H3ser10 phoshorylation (H3ser10p), a mitosis-specific histone mark, and DAPI in asynchronous and mitotic populations. (C) H3ser10p immunofluorescence analysis of asynchronous and mitotic cells. (D) FACS histograms showing the percent of cells with a DNA content of 2N (G1) and 4N (G2/M) for a representative mitotic release time course. (E) Utilizing the FUCCI2a system (Mort et al., 2014), FACS plots from a representative time course indicating the percent of cells in EG1 (Cdt1−, Gmn−), LG1 (Cdt1+, Gmn−), and S/G2/M (Gmn+) at each time point.

### Distinct waves of gene and enhancer reactivation during mitotic exit

We first investigated the transcriptional reactivation of genes and enhancers in PSCs during the mitotic release time course. We adapted the PRO-seq method for mitotic cells by permeabilizing the cells instead of isolating nuclei and verified that mitotic cells retained their purity and morphology (Figures S1D and S1E). Drosophila nuclei were used as spike-in controls to account for global transcriptional changes across time points and replicates (Table S1). We found a high correlation between biological replicates and a clear separation of mitotic samples from the rest (Figure S2A). As expected (Gottesfeld and Forbes, 1997; Konrad, 1963; Taylor, 1960), transcription was dramatically decreased during mitosis, though some genes (n=4008) retained residual expression (normalized RPKM >1) in mitotic cells (Figure 2A), in agreement with recent studies reporting partial maintenance of transcription during mitosis (Liang et al., 2015a; Liu et al., 2017a; Palozola et al., 2017).

**Figure 2.**
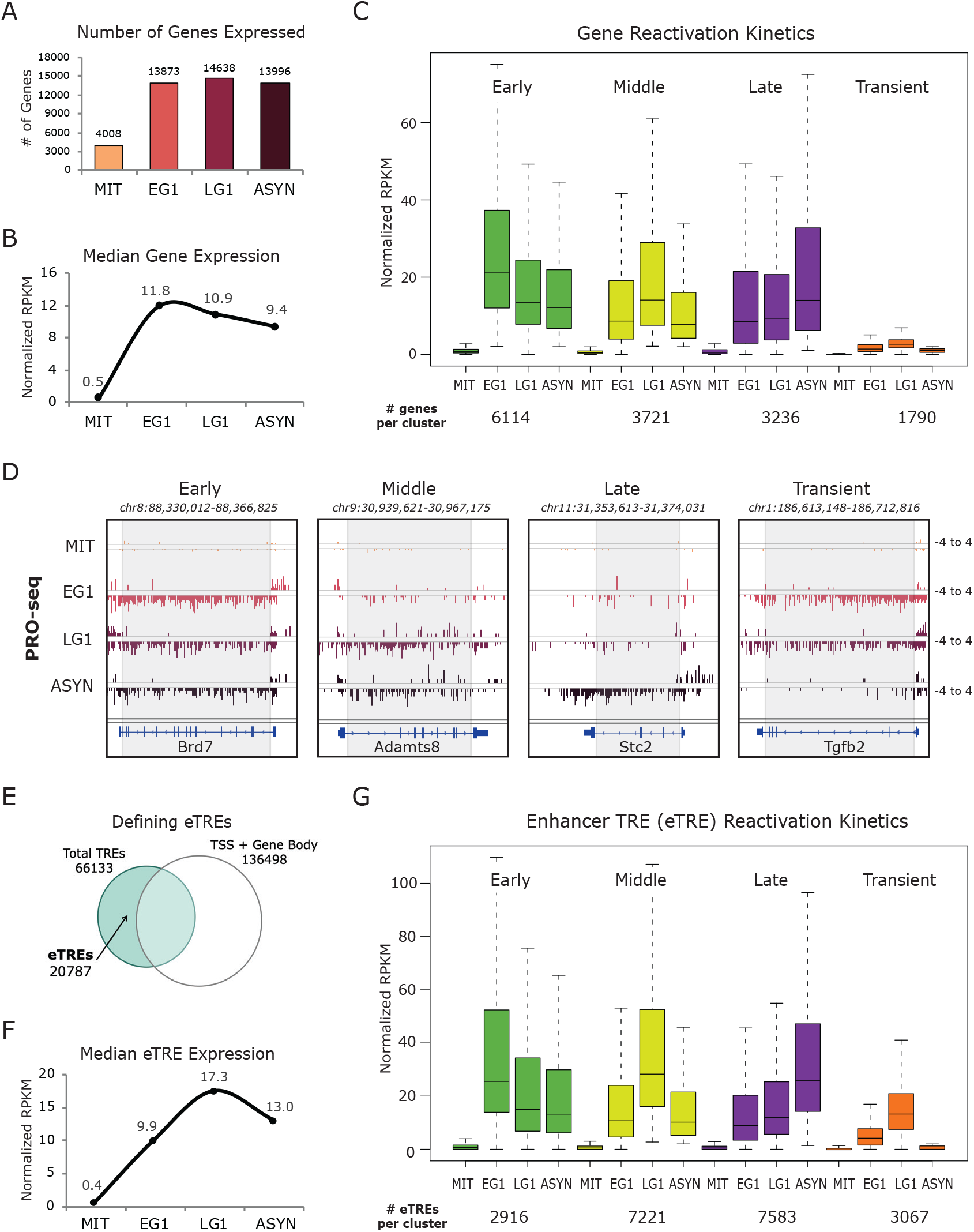
Distinct waves of gene and enhancer reactivation during mitotic exit. (A) Number of genes expressed at each time point, where expression is defined as reads per kilobase of transcript, per million mapped reads (normalized RPKM) >1, after normalization to Drosophila spike-in control and number of cells. (B) Median expression at each timepoint of all 14861 expressed genes. (C) Four patterns of transcriptional reactivation kinetics: genes were assigned to a group based on when they reached their maximal expression level (EG1 = early, LG1 = middle, ASYN = late). Transient genes were defined as normalized RPKM <2 in ASYN but >2 in EG1 and/or LG1. Box plots depict the median transcriptional activity across the time course and the number of genes in each cluster are listed below. (D) Genome browser tracks show PRO-seq data (one replicate, plus and minus strands) for one gene from each reactivation group. Grey box approximates the region used to quantify PRO-seq signal after deleting the first 500bp and last 500bp of each gene to avoid the enriched PRO-seq signal that comes from the TSS and TES regions. (E) Venn diagram representing our strategy for identifying enhancer transcriptional regulatory elements (eTREs) using PRO-seq and published TSS data. Total TREs were defined using dREG algorithm (Danko et al., 2015). (F) Median expression at each time point of all 20787 identified eTREs. (G) Four patterns of eTRE transcriptional reactivation kinetics, defined as in (C). Box plots depict the median transcriptional activity across the time course and the number of eTREs in each cluster are listed below.

Globally, transcription was rapidly reset by EG1, with similar numbers of genes expressed in early G1, late G1 and asynchronous cells (Figure 2A). Median expression levels were highest in EG1 (Figure 2B), suggesting a transcriptional spike upon mitotic exit, in agreement with observations in other cell types (Hsiung et al., 2016; Palozola et al., 2017). We noticed that the kinetics of transcriptional reactivation was variable among genes and clustered genes into different groups based on when they reached their maximal expression level: early G1 (Early), late G1 (Middle), or asynchronous (Late) (Figures 2C and 2D; Table S2). We also identified a small group of genes (n=1790) that were transiently expressed in EG1 and/or LG1 but decreased to low levels (normalized RPKM <2) in asynchronous cells (Transient group). The reactivation patterns of randomly selected genes from each group were validated by RT-qPCR analysis of nascent pre-mRNA transcripts in an independent release time course as well as in FUCCI2a (Mort et al., 2014) sorted cells (Figure S2B and S2C). Other than the Transient genes, which showed much lower overall expression in asynchronous cells both by PRO-seq (Figure 2C) and RNA-seq (Figure S2D) analysis, median transcriptional levels of the gene clusters were similar and did not directly correlate with the observed reactivation patterns. Transcriptional resetting kinetics did not reflect differences in gene length, as Late reactivated genes were the shortest, on average, and Early and Middle had similar size ranges (Figure S2E). These results document that gene reactivation during mitotic exit in PSCs is an asynchronous process that is not dependent on the absolute transcriptional levels or the rate of RNA polymerase elongation.

Next, we assessed the transcriptional reactivation of enhancers, taking advantage of the ability of PRO-seq to detect all nascent RNA transcription. We were able to generate an atlas of transcriptional regulatory elements (TREs) throughout the genome using the dREG algorithm (Danko et al., 2015). We excluded all TREs within 1kb of any TSS or gene body (for both protein-coding and non-coding genes), resulting in 20,787 high-confidence enhancer TREs (eTREs) (Figure 2E). In contrast with the global transcriptional spike of genes at early G1 (Figure 2B), eTREs showed overall a slower reactivation (Figure 2F), suggesting potentially different mechanisms of transcriptional resetting. However, using the same criteria as above we could still identify eTREs that followed Early, Middle, Late, or Transient reactivation kinetics similar to those observed in genes (Figures 2G and S2H; Table S2).

### Stem cell genes and enhancers are rapidly reactivated upon mitotic exit

To gain insight into the biological relevance of the differential reactivation kinetics of genes and eTREs, we performed Gene Ontology (GO) analysis (McLean et al., 2010) and found a large number of housekeeping and cell fate-related processes associated with distinct reactivation groups. Metabolic and signal transduction processes were enriched in the Middle gene cluster, while the Late gene group preferentially included genes involved in chromosome segregation and cell division, such as *Cdca2* and *Aurka* (Figures 3A and 3B). Both Middle and Late eTREs also associated with similar functions (Figure S3A). Other general “housekeeping” processes like transcription, RNA splicing, and protein transport were mostly associated with the Early gene group (Figure 3A). Notably, only the Early gene group enriched for terms associated with stem cell maintenance (Figure 3A) and showed a strong overrepresentation within a published ESC/iPSC gene expression signature (Papadimitriou et al., 2016), which included known regulators of pluripotency such as *Nanog*, *Sall4*, and *Klf4* (Figures 3B and 3C). Early reactivated eTREs also enriched for stem cell regulation categories (Figure S3A), and the previously-defined PSC super-enhancers (SEs) (Whyte et al., 2013) were overrepresented in this group (Figure S3B), showing much faster and stronger reactivation compared to all eTREs (Figure 3D).

**Figure 3.**
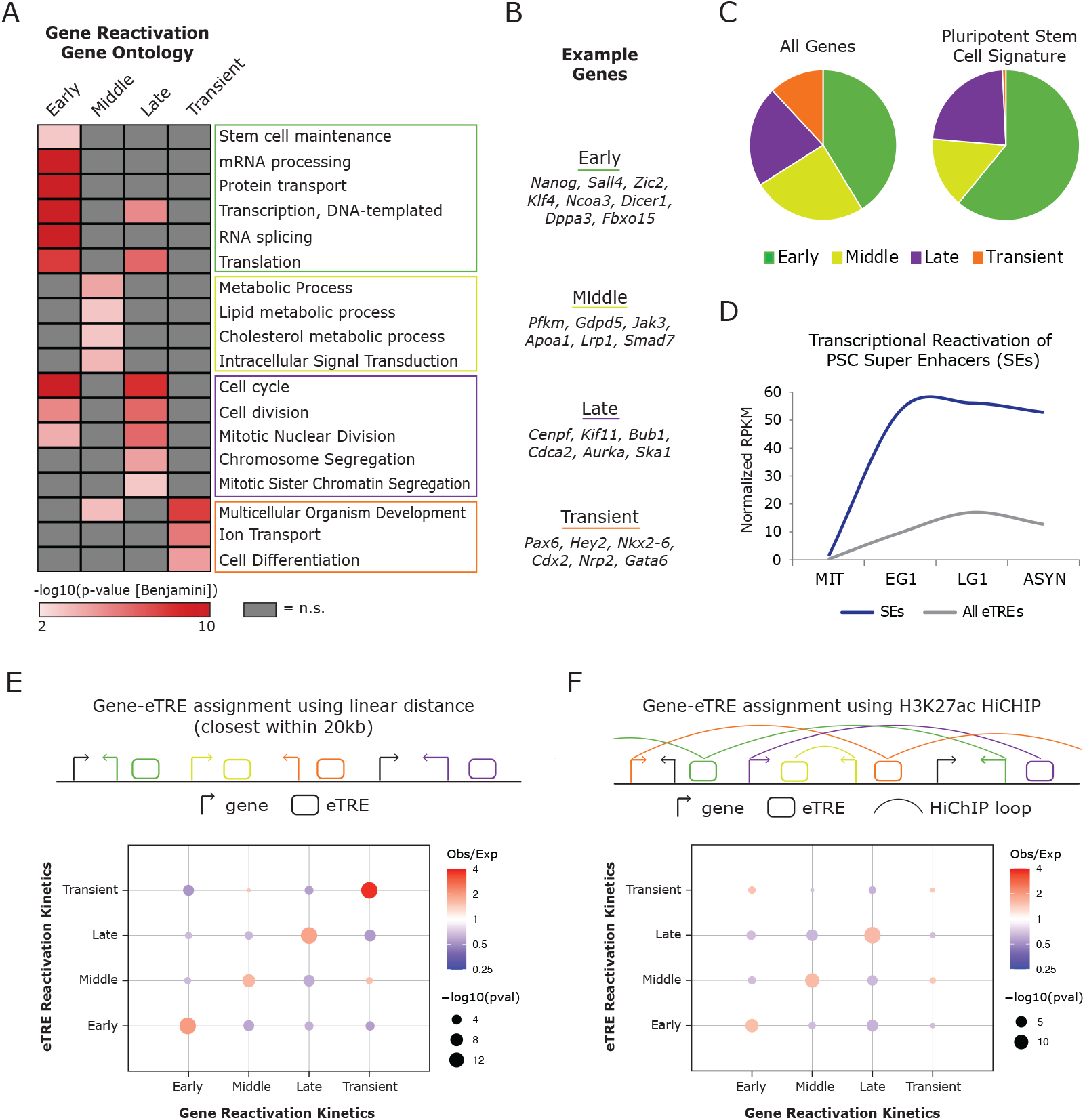
Stem cell genes and enhancers are rapidly reactivated upon mitotic exit. (A) Heat map indicates the enrichment (−log10(pvalue)) of selected top Gene Ontology (GO) terms in each gene reactivation cluster. Adjusted (Benjamini) pvalue was used with a cut-off of p<0.01. Not significant (p>0.01) terms are shown in gray. (B) Example genes in each reactivation group corresponding to top GO terms. (C) Pie charts show the distribution of all 14861 expressed genes (“All Genes”) versus 478 genes defined in an ESC/iPSC signature (Papadimitriou et al., 2016) (“Pluripotent Stem Cell Signature”) within the reactivation clusters. (D) Transcriptional reactivation of all eTREs compared to eTREs that overlap with defined PSC SEs (n=246 eTREs) (Whyte et al., 2013). See Table S6 for statistical analysis. (E) Schematic for assigning eTREs to genes based on linear distance (closest eTRE +/−20kb from TSS). Relative enrichment of transcriptional reactivation patterns of paired eTREs and genes using this method. Size of dots indicates p-value (two-sided Fisher’s exact test) and color indicates ratio of observed (Obs) versus expected (Exp) frequency. (F) Schematic for assigning eTREs to genes based on asynchronous PSC H3K27ac HiChIP data (Di Giammartino et al., 2019) at 10kb resolution. Relative enrichment of transcriptional reactivation patterns of paired eTREs and genes using this method. Size of dots indicates p-value (two-sided Fisher’s exact test) and color indicates ratio of observed (Obs) versus expected (Exp) frequency. Precise p-values and Obs/Exp ratios for all relevant panels can be found in Table S6.

Intriguingly, the transiently-activated gene and eTRE clusters enriched for developmental and differentiation processes, and included known regulators of lineage specification, such as *Gata6*, *Cdx2* and *Pax6* (Figures 3A, 3B, and S3A). In contrast with the other three gene clusters, the promoters of transient genes were not occupied by pluripotency TFs or transcriptional coactivators based on enrichment analysis with publicly available ChIP-seq data from asynchronous PSCs (Sheffield and Bock, 2016). Instead, they enriched for binding of Polycomb Repressive Complex 1 and 2 (PRC1/2) components (EZH2, SUZ12, JARID2, RING1B) (Figure S3C), further suggesting that transiently activated genes are in a more repressive/poised state during interphase. Indeed, 20% of transient genes were bivalent, based on a previously defined high-confidence bivalent gene set (Asenjo et al., 2020), as opposed to 2-10% of any other gene cluster (data not shown).

These results document distinct transcriptional reactivation modules in PSCs entering G1 that associate with the two hallmark properties of PSC identity: self-renewal and pluripotency.

### Enhancer reactivation patterns mirror the kinetics of proximal and long-range target genes

While the global transcriptional resetting patterns and the respective GO terms of both eTREs and genes were highly concordant, this does not allow drawing firm conclusions on the relative reactivation of enhancers and their target genes. To address this, we assigned eTREs to their most proximal gene based on linear distance (<20kb) (Figure 3E), and to one or more distal target genes (>20kb) (Figure 3F) based on long-range chromatin contacts detected by H3K27ac HiChIP analysis in asynchronous PSCs (Di Giammartino et al., 2019). Both approaches showed coordinated activation of eTREs and their target genes at frequencies significantly higher than expected by chance (Figures 3E and 3F). Interestingly, there were still many Early genes whose fast reactivation could not be explained by a proximal or looped Early eTRE. While this may in part be due to our strict criteria for eTRE calling, which excludes all intragenic enhancers, it also indicates that early reactivation of some genes may be promoter-driven and not dependent on enhancer activity.

### Mitotic bookmarking predicts rapid transcriptional reactivation

Despite the global perturbation of chromatin landscape and TF binding during mitosis, several of the regulatory proteins we found enriched at gene clusters in asynchronous cells (see Figure S3C) have recently been shown to be retained on mitotic chromatin at a subset of their interphase sites. These mitotic bookmarking factors include pluripotency TFs (OCT4, SOX2, KLF4, and ESRRB) (Deluz et al., 2016; Festuccia et al., 2016; Festuccia et al., 2019; Liu et al., 2017; Owens et al., 2019; Teves et al., 2016), transcriptional machinery (TBP) (Teves et al., 2018), architectural factors (CTCF) (Owens et al., 2019), as well as the active histone mark H3K27ac (Liu et al., 2017b) (Figure 4A). To investigate how the mitotic retention or loss of specific features associate with transcriptional reactivation kinetics during G1 entry, we compiled in-house and published mitotic and asynchronous ChIP-seq datasets for various bookmarking factors and also performed ATAC-seq (Buenrostro et al., 2013) in mitotic and asynchronous PSCs. For each feature, we defined peaks as “Retained” if they were present in both mitotic and asynchronous cells and “Lost” if they were only present in asynchronous cells (Figure 4B, see also methods) and then examined their enrichment around gene promoters (+/−2.5kb) and eTREs (+/−2.5kb). This analysis showed that promoters of rapidly reactivated genes (Early group) had a significantly higher likelihood to retain H3K27ac, chromatin accessibility, and TBP and KLF4 binding (odds ratio (OR) observed/expected >1.5, p<0.001) compared to all other gene groups (Figure 4C; Table S3). On the other hand, Late reactivated gene promoters only enriched for Lost H3K27ac peaks. In agreement, we observed that more than 85% of Early gene promoters overlapped with at least one Retained peak from H3K27ac, ATAC-seq, or TBP, and many (45%) retained all three features (Figure 4D). While we expect these marks to frequently co-exist in interphase cells, the retention of all three features at Early genes during mitosis supports a potentially additive effect of bookmarking. Interestingly, early reactivated eTREs correlated not only with bookmarking by H3K27ac and chromatin accessibility, but also with mitotic retention of additional pluripotency TFs, such as OCT4, SOX2, KLF4 and ESRRB (Figure S4A and S4B; Table S3). Finally, transient genes and eTREs showed no significant enrichment for any of the tested factors, highlighting the unique regulation of these regions and underscoring the need for future investigation. These findings establish a strong association between mitotic bookmarking and rapid transcriptional reactivation, propose a hierarchy in the importance of bookmarking factors for reactivation, and uncover distinct roles for certain bookmarking factors at genes versus enhancers.

**Figure 4.**
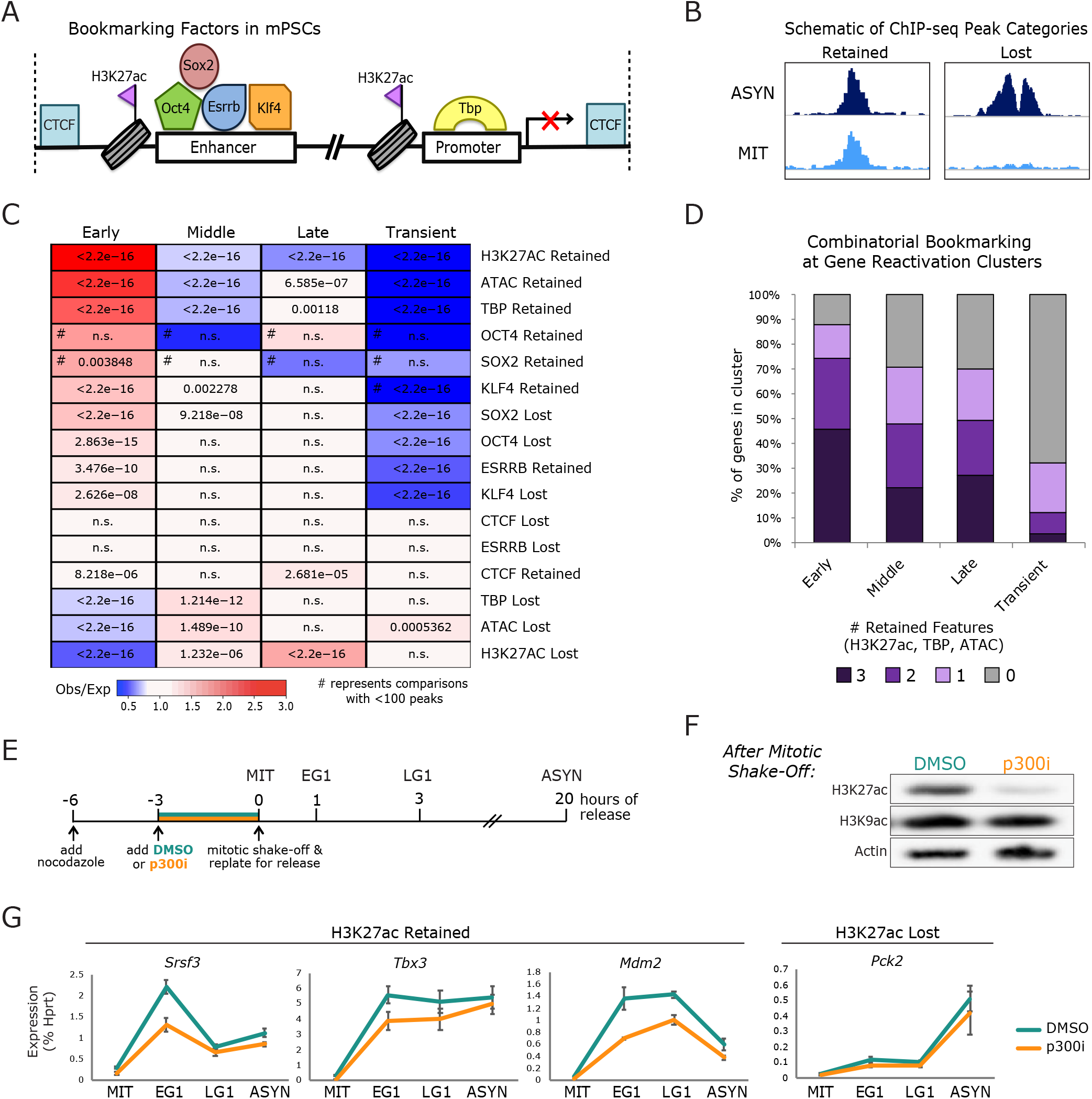
Mitotic bookmarking predicts rapid transcriptional reactivation. (A) Schematic depicting the pluripotency TFs, general TFs, architectural factors, and histone modifications that have been identified as mitotic bookmarking factors in PSCs and have available ChIP-seq both in mitotic and asynchronous cells. (B) Schematic of a Retained ChIP-seq peak (present in asynchronous and mitotic cells) and a Lost ChIP-seq peak (present only in asynchronous cells), indicating how each binding site for the bookmarking factors in (A) will be defined. (C) Relative enrichment or depletion of the Retained/Lost peaks for each bookmarking factor at the promoter (+/−2.5kb from TSS, at least 1bp overlap) of the gene reactivation clusters. Color indicates ratio of observed (Obs) versus expected (Exp) frequency and p-value (two-sided Fisher’s exact test) is indicated if significant (p<0.01). Comparisons using <100 overlapping peaks are denoted with a hash mark (#). See Table S3 for complete statistical analysis. (D) Stacked bar plot showing the percent of genes in each transcriptional reactivation cluster that retained during mitosis all three bookmarking features (H3K27ac, TBP, ATAC-seq), two of these features, one of these features, or none of them. (E) Schematic illustrating the strategy for selectively depleting H3K27ac during mitosis using a p300/CBP inhibitor (p300i) versus a DMSO-treated control. The p300i or DMSO was added three hours before mitotic shake-off. Mitotic cells were then released into fresh media without the inhibitor and were collect after 1 hour (EG1), 3 hours, (LG1), and 20 hours (ASYN) (F) Western blot analysis of mitotic populations treated for three hours with p300i or DMSO. Actin was used as a loading control and H3K9ac to show the selectivity of the inhibitor for H3K27ac and not global acetylation. (G) Transcriptional reactivation kinetics for three randomly selected bookmarked genes (“H3K27ac Retained”) and one non-bookmarked genes (“H3K27ac Lost”) after treatment with p300i or DMSO, as shown in Figure 4E. pre-mRNA qPCR from two independent mitotic release time courses is shown, error bars show +/−SEM.

To functionally test the role of H3K27ac bookmarking in rapid transcriptional reactivation, we utilized a selective p300/CBP inhibitor (p300i) (Lasko et al., 2017) to deplete H3K27ac during mitosis. Cells were treated with p300i or DMSO for three hours prior to mitotic shake-off and then released in full medium without p300i or Nocodazole for 1hr (EG1), 3hr (LG1) or 20hrs to allow full resetting (ASYN) (Figure 4E). Western blot analysis validated the specific loss of H3K27ac in the mitotic population upon p300i treatment (Figure 4F), while FACS analysis of DNA content at each time point showed no effects of the treatment on mitotic purity or cell cycle release (data not shown). Pre-mRNA qPCR analysis for three randomly selected bookmarked genes showed that p300i-treatment significantly compromised their transcriptional reactivation at EG1, although they efficiently recovered at later stages (Figure 4G). The reactivation pattern of a non-bookmarked gene remained unaffected. These data support a functional role for H3K27ac mitotic bookmarking in early transcriptional resetting.

### Chromosomal compartments and domain boundaries are established in EG1 in coordination with transcriptional reactivation

Next, we assessed the possibility that rebuilding of the 3D chromatin architecture during mitotic exit is instructive for transcriptional resetting within local domains or long-range chromatin interactions. To do so, we first performed *in situ* Hi-C in the same mitotic release time course as for PRO-seq (Figure 1A). Principle Component Analysis (PCA) confirmed consistency between our Hi-C replicates and indicated gradual changes in the course of mitotic release (Figure S5A). Consistent with previous studies (Abramo et al., 2019; Gibcus et al., 2018; Nagano et al., 2017; Naumova et al., 2013; Zhang et al., 2019), we observed a dramatic alteration of chromatin architecture during mitosis (Figure S5B) with an almost complete loss of compartmentalization, topologically-associating domains (TADs), and specific, long-range interactions (Figure 5A).

**Figure 5.**
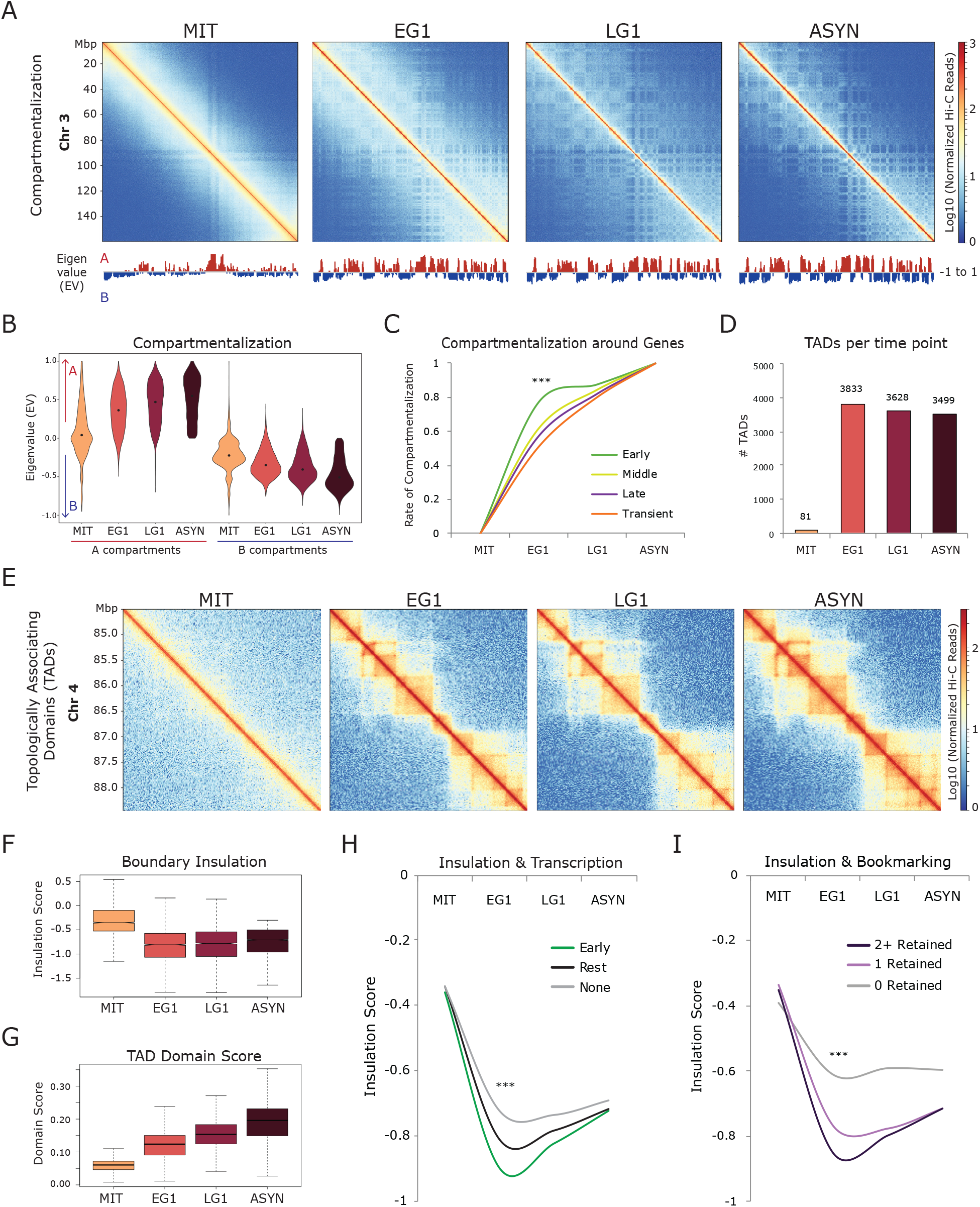
Chromosomal compartments and domain boundaries are established in EG1 in coordination with transcriptional reactivation. (A) Hi-C interaction heatmaps (log10 Normalized Hi-C reads) of chromosome 3 for each time point to illustrate compartment reformation. The eigenvalues (EV) of each matrix at 100kb resolution are shown below. (B) Violin plot depicting the compartmentalization (eigenvalue, EV) for all 100kb bins across the genome at each time point, separated by A or B compartments. Bins with a final (in asynchronous cells) EV>0 are called A, while a final EV<0 are called B. (C) Line plot showing the rate of compartmentalization during mitotic exit for bins containing genes from each of the four reactivation kinetic clusters (Early, Middle, Late, Transient). If a bin contained genes from multiple clusters, it was prioritized as Early (n=4047) > Middle (n=2901) > Late (n=2278) > Transient (n=1636). *** indicates p<0.0001 for Early versus any other cluster, two-sided Wilcoxon’s rank sum test. See Table S6 for complete statistical analysis. (D) Number of TADs identified at each time point. (E) Hi-C interaction maps (log10 Normalized Hi-C reads) of a region on chromosome 4 (chr4:84,500,000-88,500,000) for each time point. (F)Box plot showing the insulation score for the boundaries of asynchronous TADs during the time course. (G) Box plot showing the domain score for asynchronous TADs during the time course. (H) Median insulation score at each time point for TAD boundaries containing: at least one early gene or eTRE (Early, n=865), only later activated genes or eTREs (Middle, Late, and/or Transient) (Rest, n=1171), or no active genes or eTREs (None, n=1483). Only TAD boundaries called in asynchronous cells were used, and *** indicates p<0.0001 for all pairwise comparisons, two-sided Wilcoxon’s rank sum test. See Table S6 for complete statistical analysis. (I) Median insulation score at each time point for TAD boundaries that retain during mitosis: 2+ bookmarking features (CTCF, H3K27ac, ATAC-seq, TBP), one bookmarking feature, or none. *** indicates p<0.0001 for all pairwise comparisons, two-sided Wilcoxon’s rank sum test. See Table S6 for complete statistical analysis.

Upon mitotic exit, the majority (~85%) of A and B compartments (Lieberman-Aiden et al., 2009) (defined as 100kb bins with eigenvalue (EV)>0.2 or <−0.2, see also methods) (Figure S5C), were reestablished by EG1, though they continued to segregate and gain strength at later time points (Figures 5A and 5B). The vast majority of expressed genes regardless of their reactivation kinetics were located within A compartments (defined as open, gene-rich and active genomic regions), with the exception of 462 transiently expressed genes, which resided within B compartments (Figure S5D). Although absolute compartmentalization scores were similar between early, middle, and late reactivated genes in asynchronous cells (Figure S5E), compartments that harbored early reactivated genes (and eTREs) reached their final compartmentalization strengths at a significantly faster rate than ones with gradually or transiently reactivated genes (Figures 5C and S5F). These findings show that the kinetics of transcriptional resetting correlate with the recovery rate of chromosomal compartments.

3D chromatin organization at the level of TADs was also disrupted in mitosis and largely reset by EG1 (Figures 5D and 5E), in agreement with previous reports in PSCs and other cell types (Abramo et al., 2019; Nagano et al., 2017; Zhang et al., 2019). Boundary insulation was rapidly recovered (Figure 5F), while TAD domain score, which measures the ratio of intra-TAD versus overall connectivity (Dixon et al., 2012; Stadhouders et al., 2018), increased more gradually during G1 entry (Figure 5G). A substantial fraction (~20%) of asynchronous TADs were formed by the merging of smaller TADs in EG1 and/or LG1 (data not shown), supporting a recently proposed bottom-up model of TAD formation (Zhang et al., 2019). Interestingly, the rate of transcriptional reactivation of TAD-associated genes, although not well correlated with the resetting of the domain score, showed a significant association with TAD boundary insulation. Boundaries that overlapped with at least one Early gene/eTRE showed significantly faster and stronger insulation compared to boundaries with later reactivated genes/eTREs or with no transcriptional activity at any stage (Figure 5H). Moreover, boundaries with stronger insulation scores at EG1 compared to asynchronous (“Early Spike”) were enriched for early-reactivated genes, supporting a link between the observed transcriptional and insulation spikes at EG1 (Figure S5G). On the other hand, gradually-insulated boundaries (“Gradual”) showed no preference for any gene reactivation group. Together, these results establish strong links between the timing and degree of insulation with transcriptional activity and resetting upon mitotic exit.

Given that proteins with known or presumed architectural roles, such as CTCF or transcription factors, have been reported as bookmarking factors in PSCs (Liu et al., 2017b; Owens et al., 2019; Teves et al., 2018), we tested whether mitotic retention of histone marks and/or chromatin-bound proteins could facilitate faster architectural resetting during mitotic exit. Indeed, boundaries that retained H3K27ac, CTCF, or other bookmarking features (TBP and ATAC-seq) during mitosis established significantly stronger insulation by EG1 compared to the ones that lost these features (Figure S5H-K). Mitotic retention of multiple (2+) factors and marks predicted even faster and stronger insulation during G1 entry (Figure 5I), suggesting additive effects. The potential significance of mitotic bookmarking for topological resetting is further underscored by the observation that 95% of TAD boundaries (3343/3519) were bookmarked by at least one factor (H3K27ac, TBP, CTCF, and/or ATAC-seq).

Together, these findings show a rapid resetting of chromosomal compartments and TAD boundaries that correlates with transcriptional reactivation and mitotic bookmarking.

### Active regulatory elements and bookmarked regions engage rapidly in chromatin looping, while structural loops reform at a slower rate

Next, we assessed the kinetics of chromatin loops during mitotic exit. Significant contacts were identified by applying Fit-Hi-C (Ay et al., 2014) at 20kb resolution with a cut-off of q-value<10-3 for both replicates per time point. Similar to the progressive increase in TAD domain score (see Figure 5G), loops appeared to reform in a gradual manner (Figure S6A). K-means clustering of all 52,489 high-confidence contacts across all time points identified three groups of interactions with different kinetics of resetting: Fast, Gradual, and Slow (Figures 6A and 6B; Table S4). All groups enriched for binding of the classical architectural factors CTCF and cohesin (Figure 6C) and showed a progressive increase in size (Figure S6B), suggesting that more distal contacts are formed slower than closer ones in agreement with the loop extrusion model (Fudenberg et al., 2016; Ganji et al., 2018; Guo et al., 2015; Nora et al., 2017; Rao et al., 2017; Sanborn et al., 2015). Association analysis revealed that Fast established contacts likely represent enhancer-promoter interactions, since their anchors were strongly enriched for active histone marks, binding of pluripotency TFs, and transcriptional machinery (Figure 6C). Gradual and Slow contacts showed a stronger enrichment for CTCF and Cohesin binding and no association with transcriptional regulatory features, indicating that CTCF-Cohesin structural loops that do not overlap with enhancers and promoters are formed at a slower rate during mitotic exit. We also observed a significant -but moderate- enrichment for PRC1/2 components (RNF2, EZH2, SUZ12, JARID2), particularly at Slow contacts, suggesting that repressive PRC-mediated contacts may be reestablished later in G1 (Figure 6C). Examples of differential loop kinetics are presented by Virtual 4C plots (Figure S6C) and Hi-C contact maps (Figure S6D).

**Figure 6.**
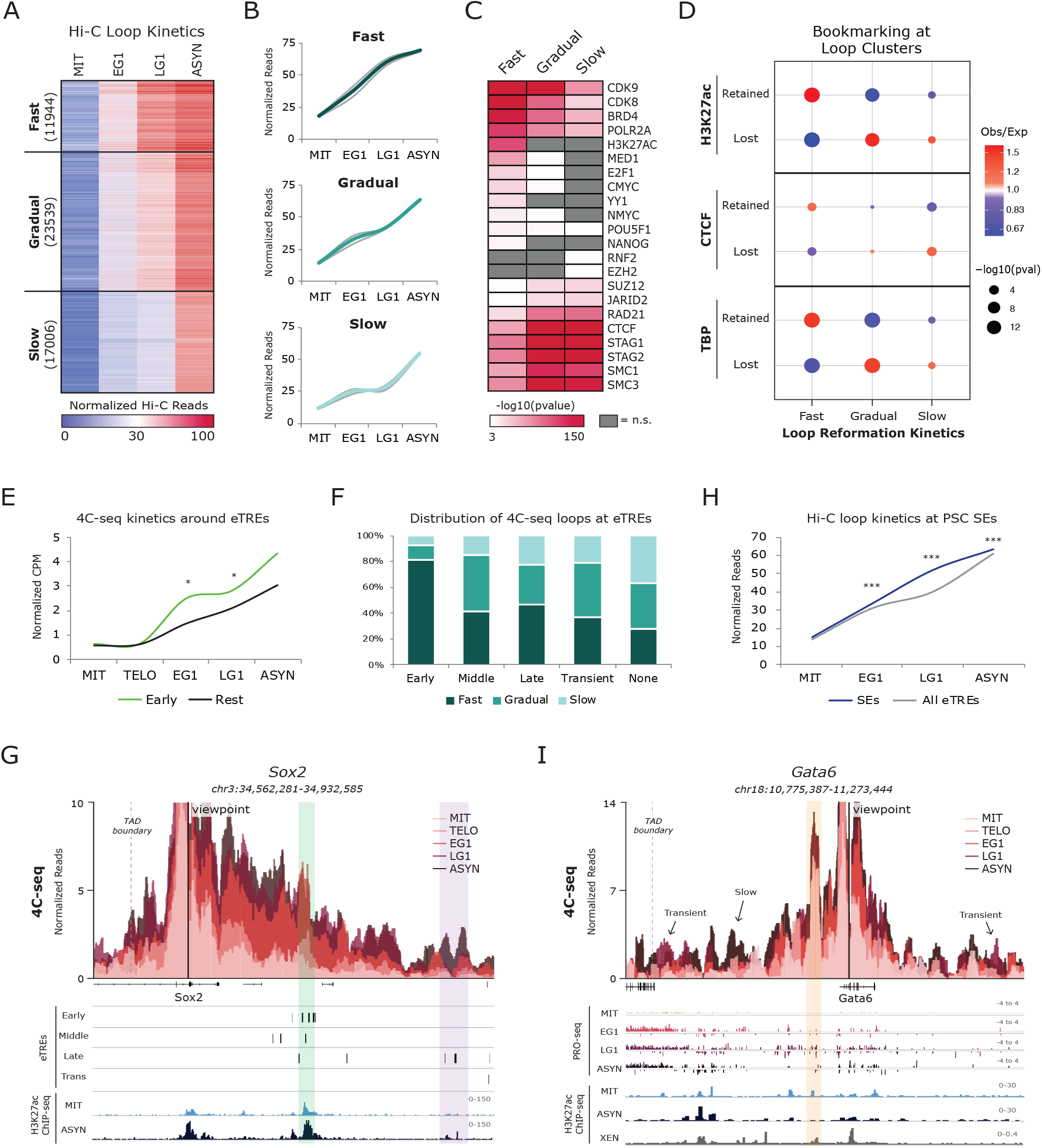
Enhancer-promoter contacts are rapidly reformed with different patterns at early or transiently activated genes. (A-B) K-means clustering of all 52489 Hi-C loops. (A) Heat map of the three reformation clusters (Fast, Gradual, Slow) illustrating the normalized Hi-C reads over the time course for each contact. Number of loops per cluster is shown. (B) Line plots depicting the median of normalized read counts for each cluster over the time course. Median of each replicate is depicted with gray lines while the average of the replicates is shown in color. (C) Heat map indicates the enrichment (−log10(pvalue)) of select factors binding in the accessible regions of the anchors for each Hi-C loop cluster as calculated from LOLA analysis (Sheffield and Bock, 2016). Not significant (q>0.001) terms are shown in gray. (D) Relative enrichment of Hi-C loop reformation clusters with presence of a Retained/Lost H3K27ac, CTCF, or TBP ChIP-seq peak. Size of dots indicates p-value (two-sided Fisher’s exact test) and color indicates ratio of observed (Obs) versus expected (Exp) frequency. Anchors in multiple loop clusters were prioritized by Fast > Gradual > Slow. Anchors overlapping multiple peaks were prioritized by Retained > Lost. (E) Line plot showing the 4C-seq contact strength over the time course for loop anchors that overlap with or are nearby to (<5kb) Early eTREs (Early, n=17 loops) versus Middle, Late, or Transient eTREs (Rest, n=48 loops). Loop anchors <5kb from multiple eTREs were prioritized as Early>Rest. * indicates p<0.05 for Early vs. Rest, two-sided Wilcoxon’s rank sum test. See Table S6 for complete statistical analysis. (F) Stacked bar plot showing the percent of 4C-seq Fast, Gradual, and Slow reformed loops overlapping or near to (<5kb) Early (n=27), Middle (n=46), Late (n=49), or Transient (n=19) eTREs. Any loop anchor >5kb from all eTREs was shown as “None” (n=167). Loop anchors <5kb from multiple eTREs were counted multiple times, once for each eTRE. (G) 4C-seq data is represented as average CPM around the viewpoint (*Sox2* promoter) with each time point shown as an overlapping bar plot. Genome browser tracks underneath show eTRE reactivation clusters (Early, Middle, Late, and Transient) and H3K27ac ChIP-seq in mitotic and asynchronous cells. The Sox2 super-enhancer (Early eTREs) is highlighted in green while another region (Late eTREs) is highlighted in purple. (H) Line plot showing Hi-C contact strength for loops with an anchor containing a PSC SE (Whyte et al., 2013) (n=813) versus loops containing any eTREs (n=19559). *** indicates p<0.0001 for SEs vs. All eTREs, two-sided Wilcoxon’s rank sum test. See Table S6 for complete statistical analysis. (I) 4C-seq data is represented as average CPM around the viewpoint (*Gata6* promoter) with each time point shown as an overlapping bar plot. Genome browser tracks underneath show raw PRO-seq reads for each time point, H3K27ac ChIP-seq in mitotic and asynchronous cells, and H3K27ac ChIP-seq in asynchronous eXtraembryonic ENdoderm (XEN) stem cells. A retained contact between the Gata6 promoter and a proximal enhancer is highlighted in orange. Arrows indicate visually-detected transient G1 loops and a slow-formed contact.

We next examined the role of mitotic bookmarking in loop reformation kinetics. Mitotic retention of TBP or H3K27ac enriched at the anchors of fast-established contacts while loss of these factors correlated with gradual or slow loop formation (Figure 6D). Mitotic retention or loss of CTCF showed similar, but less significant, associations. This analysis suggests that -in addition to promoting rapid transcriptional resetting-mitotic bookmarking by H3K27ac and/or TBP is also predictive of faster loop reformation.

### Different patterns of chromatin loops characterize early or transiently activated genes and enhancers that promote self-renewal or differentiation

The coordinated reactivation of enhancers and their target genes during mitotic exit (see Figures 3E and 3F) suggests a functional link between chromatin looping and transcriptional reactivation. This is further emphasized by the correlation of both looping and transcriptional resetting with the retention of certain mitotic bookmarking factors. We first examined the temporal relationship between transcriptional reactivation and fine-scale architectural resetting using the Hi-C loop clusters. Genes and eTREs within early reformed contacts had significantly higher transcriptional activity compared to the other clusters (Figure S6E), in agreement with a regulatory nature of these loops as indicated by the enrichment analysis of the anchors. However, transcriptional recovery rates were indistinguishable across loop clusters. We considered this may be due to the technical limitations of Hi-C analysis, which misses short-range interactions (<60kb) (see Methods), enriches for structural over regulatory loops (Bonev et al., 2017; Di Giammartino et al., 2019; Mumbach et al., 2016), and does not cover a large fraction (~50%) of expressed genes and eTREs. Therefore, we decided to revisit this question by performing high-resolution 4C-seq around the promoters of 11 example genes with distinct reactivation patterns and biological relevance (Table S5). In addition to the previous mitotic release time course, we also included an earlier time point (30 minutes release, see Figure S1B) in order to capture chromatin contacts during anaphase/telophase (TELO), as reported in other cell types (Zhang et al., 2019).

For each viewpoint, we called significant contacts at 500bp windows based on differential normalized 4C-seq signal between any two time points (Fold change >3, p<0.05). After merging adjacent bins (<1kb) and filtering out loops with median strength <2 in asynchronous cells, we detected a total of 232 high-confidence contacts, which showed gradual and nonsynchronous reformation kinetics (Figure S6F; Table S4), as observed in Hi-C. While proximal loops were often among the first reactivated, many loci showed clear exceptions to this rule, with more distal loops reaching their maximal strength prior to proximal contacts (Figure S6G). This suggests that while distance is a limiting factor in the rate of loop resetting, additional features, such as local chromatin features and transcriptional kinetics might also play a role. Indeed, we found that loops connecting genes to Early eTREs were significantly faster reformed than those contacting later (Middle, Late, Transient) eTREs or no eTREs (Figures 6E and 6F). For example, at the *Sox2* locus (a critical regulator of pluripotency and reprogramming) we observed rapid contact reformation between the promoter and super enhancer (green highlight) (Whyte et al., 2013), which contains multiple Early eTREs, and much slower interaction with nearby Late eTREs (purple highlight) (Figure 6G). This was true for all rapidly reactivated genes that we tested, which formed at least one fast loop with an Early eTRE prior to other contacts (see another example in Figure S6H). This suggests that this early enhancer-promoter communication is sufficient for initial gene activation, while later loops may stabilize gene expression or provide additional layers of regulation. Reanalysis of our Hi-C data independently confirmed the faster reformation of contacts around pluripotent stem cell super-enhancers (Whyte et al., 2013) (Figure 6H). These results support that rapid architectural reorganization during mitotic exit is linked to fast transcriptional reactivation, particularly around stem cell-related features.

Although the vast majority of observed 4C-seq contacts were abrogated in mitosis and reestablished only at EG1 or later stages, we observed several noteworthy exceptions. For example, when we focused on the transiently expressed *Gata6*, a known master regulator of the endodermal fate (Schrode et al., 2014), we observed a strong, proximal contact (orange highlight) that was maintained in mitosis and G1 phase (Figure 6I). This anchor corresponded to a putative enhancer in eXtra-embryonic ENdoderm (XEN) cells (unpublished H3K27ac ChIP-seq from our group), implying that this persistent contact may pre-program activation of *Gata6* during G1 and/or upon endoderm differentiation. We also discovered transiently formed loops during G1, which differ from the recently reported “aberrant” inter-TAD chromatin contacts that are disrupted upon the late establishment of intervening boundaries (Zhang et al., 2019). The observed transient contacts occurred within the same TAD and coincided with the transient activation of genes or eTREs at their anchors, suggesting a potentially activating role. Finally, we detected several late-formed loops around Gata6 and other transient viewpoints (Figures 6I and S6I), which could act to repress lineage-specific genes and enhancers after G1 as also indicated by the Hi-C analysis (Figures 6C and 6D). Future functional characterization of these loops will shed light on their role during the balance of self-renewal and differentiation in stem cells during G1.

Taken together, our analyses during mitotic exit revealed complex patterns of regulatory contacts that associate with the transcriptional reactivation kinetics of involved elements and may thus contribute to the proper maintenance of PSC identity.

## DISCUSSION

The MIT-to-G1 transition poses substantial challenges for the maintenance of cell identity. To better understand how cells achieve molecular and functional resetting of their identity after mitosis, we have generated the first integrative map of transcriptional and architectural changes during mitotic exit in PSCs. Our analyses revealed that various molecular characteristics are reset in a nonsynchronous manner and provided insights into the potential factors and features that might determine the kinetics and efficiency of transcriptional resetting (see model in Figure 7). In particular, our data support a regulatory function of mitotic bookmarking for the rapid reestablishment of cell type-specific architectural features and transcriptional programs after cell division.

**Figure 7.**
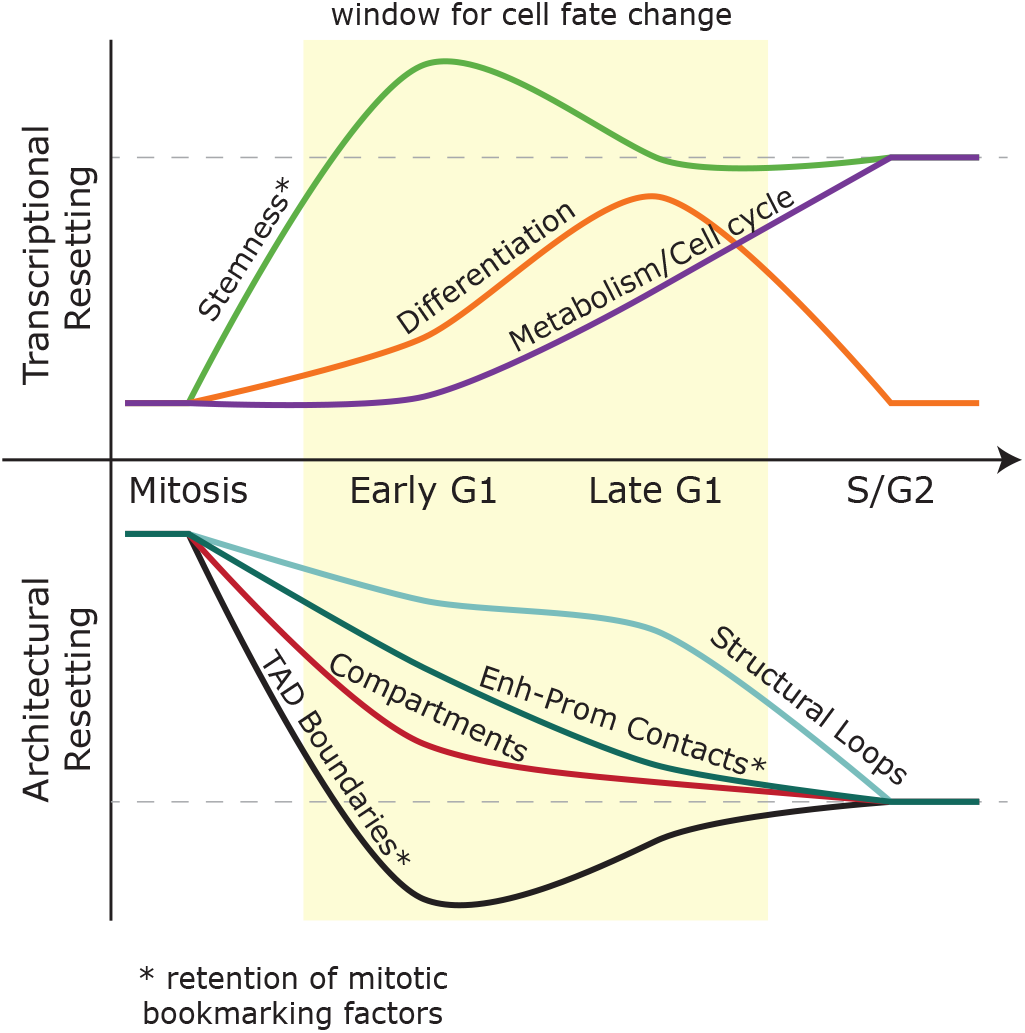
Resetting of PSC identity during G1 entry. Model summarizing the transcriptional and topological resetting of pluripotent stem cell identity during G1 entry. The top half describes the waves of transcriptional (re)activation while the bottom half shows the kinetics of 3D chromatin architecture reorganization. In both cases, the dashed line indicates full resetting of the interphase state.

As has been previously described (Hsiung et al., 2016; Palozola et al., 2017; Teves et al., 2018), we observed a dramatic decrease in transcriptional activity in mitotic cells, a global spike in transcription during G1 entry, and distinct waves of transcriptional reactivation. In marked contrast to somatic cells (Palozola et al., 2017), our analysis showed that genes and enhancers involved in regulation of stem cell identity undergo rapid reactivation, while genes involved in housekeeping functions (metabolism, cell cycle) were turned on in a more gradual fashion. This suggests cell type-specific differences in the transcriptional resetting upon mitotic exit and a preference of PSCs for early activation of cell identity genes. This may reflect the constant balance between self-renewal and differentiation in fast-cycling PSCs (Evans, 2011) and the particularly critical role of G1 phase in this decision (Boward et al., 2016). As some pluripotency master regulators have been shown to be degraded during mitosis, such as NANOG (Liu et al., 2017b), this rapid reactivation of stemness genes may be critical for stem cell self-renewal. On the other hand, the observed transient activation of developmental genes may constitute a temporal “priming” step that enables PSCs to respond to differentiation cues during G1 and tip the balance from self-renewal to lineage specification (Coronado et al., 2013; Pauklin et al., 2016; Pauklin and Vallier, 2013; Singh et al., 2013; Singh et al., 2015). This is in agreement with previous reports of “noisy” de-repression of lineage regulators during G1 in self-renewing human (Singh et al., 2013) and mouse (Asenjo et al., 2020) PSCs. Future single-cell and functional analyses will be required to determine the transcriptional levels and degree of co-expression of different developmental regulators in PSCs exiting mitosis and the significance of these transcriptional behaviors for pluripotency.

Our study also revealed distinct and nonsynchronous patterns of topological reorganization in PSC during mitotic exit and provided insights into their underlying mechanisms. We found that compartmentalization and TAD boundary insulation were rapidly reestablished in early G1, in agreement with other studies (Abramo et al., 2019; Dileep et al., 2015; Nagano et al., 2017; Naumova et al., 2013; Zhang et al., 2019), while long-range contacts were formed overall at a slower rate. Most of the Hi-C detected chromatin contacts were bound by CTCF and cohesin and their size was inversely correlated with the rate of recovery, suggesting that longer loops require more time to reform. This supports that CTCF/cohesin-mediated loop extrusion (Fudenberg et al., 2016; Sanborn et al., 2015) is a major mechanism of chromatin reorganization upon mitotic exit, as recently described in other cell types (Abramo et al., 2019; Zhang et al., 2019). However, both our Hi-C and 4C-seq analyses revealed that active regulatory loops marked by H3K27ac and binding of TFs and other cofactors are reset at a faster rate than structural CTCF/cohesin loops that do not involve enhancers or promoters. The resetting of regulatory prior to structural contacts, which was also reported in a recent study in erythroblasts (Zhang et al., 2019), supports that additional forces besides loop extrusion are involved in the architectural reorganization upon mitotic exit. Affinity or segregation through homotypic histone modifications and recruitment of TFs/cofactors that mediate contact formation via dimerization or phase separation are some of the potential underlying mechanisms (Kim and Shendure, 2019; Nuebler et al., 2018; Rada-Iglesias et al., 2018; Stadhouders et al., 2019).

The observed coordination between rapid architectural reorganization and transcriptional reactivation suggests tight links between the two processes and/or common mechanisms dictating their resetting. Mitotic bookmarking by protein factors or histone marks has been previously proposed to promote rapid transcriptional resetting of cell identity genes during G1 entry (Kadauke and Blobel, 2013; Palozola et al., 2019; Zaidi et al., 2010). However, direct links between mitotic retention and reactivation kinetics are rather sparse and contradictory (Behera et al., 2019; Blobel et al., 2009; Caravaca et al., 2013; Deluz et al., 2017; Dey et al., 2009; Festuccia et al., 2016; Hsiung et al., 2016; Kadauke et al., 2012; Oh et al., 2020; Owens et al., 2019; Teves et al., 2018). While some of these disparities could be due to cell type or the bookmarking factor itself, many of the previous studies tested only a limited number of genes at a time or measured global RNA levels rather than nascent transcription. By integrating our PRO-seq results that track transcriptional activity at single base-pair resolution with ChIP-seq data of mitotic bookmarking factors in mouse PSCs, we found a strong correlation between mitotic retention of H3K27ac and selected DNA-bound factors, such as TBP for genes and ESRRB/SOX2 for enhancers, and rapid transcriptional resetting. Importantly, mitotic retention-especially of H3K27ac and TBP- was also associated with faster insulation and loop formation, establishing a previously unappreciated link between bookmarking and chromatin reorganization. On the other hand, loss of these marks/ factors during mitosis associated with a slower transcriptional and architectural resetting. This suggests that “de novo” re-assembly of a favorable chromatin state and/or transcriptional complex during G1 entry are rate-limiting factors. Importantly, we functionally validated the bookmarking function of our top (most predictive) candidate, H3K27ac, which is also retained during mitosis in other cell types (Behera et al., 2019; Hsiung et al., 2016; Javasky et al., 2018; Liu et al., 2017a). Given that mitotic retention of H3K27ac is prominent around cell-type defining genes and enhancers (Liu et al., 2017b), our results support an important role of H3K27ac bookmarking both in the transcriptional and topological resetting of cell identity after division, and urge for future investigation of the mechanisms that preserve this mark during mitosis.

In conclusion, we observed a heterogenous yet coordinated resetting of transcription and 3D chromatin architecture during mitotic exit which prioritizes active, stem cell identity features and is broadly predicted by the retention of specific bookmarking factors. These findings provide insight into the order of molecular events during G1 entry in PSCs and constitute a stepping stone towards understanding the mechanisms ensuring the coordination of transcriptional and architectural resetting and ultimately the maintenance of stem cell identity.

## ACKNOWLEDGEMENTS

We are grateful to Matthias Stadtfeld, Ari Melnick, Dan Landau, Chris Barbieri, Danwei Huangfu, Steve Josefowicz, Kat Hadjantonakis, Todd Evans, and members of the Apostolou, Tsirigos, Danko and Stadtfeld laboratories for critical reading of the manuscript. We are thankful for advice from Charles Danko for dREG analysis, Boaz Aronson for 4C-seq primer design, and Laurianne Scourzic for FACS analysis and sorting. Finally, the R26Fucci2aR mice were a generous gift from the Hadjantonakis lab.This work was supported by the NIH Director’s New Innovator Award (DP2DA043813) and the Tri-Institutional Stem Cell Initiative by the Starr Foundation. BPW is supported by the NICHD with a T32 (T32HD060600) and an F30 (F30HD097926), as well as by a Medical Scientist Training Program grant from the NIGMS under award number T32GM007739 to the Weill Cornell/Rockefeller/Sloan Kettering Tri-Institutional MD-PhD Program. A.T. is supported by the American Cancer Society (RSG-15-189-01-RMC), the Leukemia and Lymphoma Society and the St. Baldrick’s Foundation.

## AUTHOR CONTRIBUTIONS

EA and BPW conceived and designed the study and wrote the manuscript together with AP and help from all authors. BPW performed all experiments with help from JL and DCDG. AP analyzed all Hi-C, RNA-seq, ATAC-seq, 4C-seq, and PRO-seq results and performed every part of the integrative computational analysis under the guidance of AT. AK assisted with the Hi-C analysis and visualization. LW performed PRO-seq experiments under the supervision of LC, who also provided input on the PRO-seq analysis. EA supervised the whole study and analyses.

## COMPETING INTERESTS

The authors declare no competing financial interests.

## METHODS

### LEAD CONTACT AND MATERIALS AVAILABILITY

Further information and requests for resources and reagents should be directed to and will be fulfilled by the Lead Contact, Effie Apostolou. This study did not generate new unique reagents.

### EXPERIMENTAL MODEL AND SUBJECT DETAILS

#### Cell line generation

Mouse embryonic fibroblasts (MEFs) were isolated from E13.5 embryos from *R26Fucci2aR* (Mort et al., 2014) mice in a mixed C57BL/6, DBA/2, and CD1 background (gift from Hadjantonakis lab (MSKCC)). Male MEFs were co-infected with a FUGW-rtTA (Maherali et al., 2008) and TRE-OKSM (STEMCCA) cassette (Sommer et al., 2009) according to standard protocols (Di Giammartino et al., 2019) and were then reprogrammed in the presence of 1μg/ml of doxycycline and 50μg/ml of ascorbic acid for 12 days prior to dox withdrawal for establishment of transgene-free induced PSCs (iPSCs). Individual iPSC clones were picked and screened for successful reprogramming based on expression of multiple pluripotency markers.

### METHOD DETAILS

#### Culture conditions

Two independent, male iPSC clones (described above) were used as biological replicates for all analyses in this study. iPSCs were cultured at 37°C on irradiated feeder cells in FBS/LIF+2i conditions. Specifically, KO-DMEM media (Invitrogen) supplemented with 15% heat-inactivated fetal bovine serum, GlutaMAX, penicillin-streptomycin, non-essential amino acids, beta-mercaptoethanol, and 1000 U/ml LIF, plus 2i (1uM MEK inhibitor (Stemgent 04-0006) and 3uM GSK3 inhibitor (Stemgent 04-0004-10)) was used.

#### Mitotic arrest, release, and cell cycle analysis

To arrest cells in mitosis, 80-90% confluent mouse iPSCs were passaged 1:3 on gelatinized plates the day before synchronization. Nocodazole (200ng/ ml) (Sigma, M1404) was added to the medium for 6hr prior to collection by mitotic shake-off. 1/3 of cells were processed immediately (for mitotic time point), while the remaining cells were washed 2x with PBS and released into pre-warmed (37°C) media and plated on gelatin. Cells were then collected at 1hr and 3hrs after release. Aliquots from each time point were used for estimation of synchronization and release efficiency by FACS (BD FACS Canto II analyzer and BD FACS Diva v8.0.3 software) after ethanol fixation and staining with 300nM DAPI (Biolegend, 422801). Aliquots from each time point were also tested for their cell cycle stage by using a BD FACS Aria II to assess the FUCCI2a markers (Cdt1-mCherry, Gmn-mVenus). Any mitotic shake-off experiment with <95% 4N cells was discarded. Asynchronous cells were processed identically but without Nocodazole treatment. FlowJo (v10.6.1) software was utilized for analysis and representation of FACS data.

#### Mitotic arrest and release with p300 inhibitor

Cells were prepared for synchronization as above. After 3 hours of Nocodazole treatment, either 10uM p300/CBP catalytic inhibitor (A485, Tocris #6387) or the same volume DMSO was added to the media for the remaining 3 hours of synchronization. After mitotic shake-off, 1/4 of cells were processed immediately (mitotic time point) while the remainder were washed and released for collection at 1hr, 3hrs, and 20hrs. At each time point, cells were processed for pre-mRNA RT-qPCR, Western Blot, and cell cycle analysis. Cell cycle analysis was performed as described above.

#### Validation of mitotic purity

The efficiency of synchronization and mitotic shake-off was estimated by FACS after ethanol fixation and staining with H3Ser10p antibody (Abcam, ab47297) and DAPI. Mitotic purity was further validated after the 1% formaldehyde fixation step of Hi-C (see below). Aliquots of mitotic and asynchronous cells were removed after fixation and 60K cells were cytospun at 1000rpm for 5mins onto a slide, following by immunofluorescence staining with H3Ser10p and DAPI. For validation of mitotic purity after PRO-seq permeabilization, aliquots of mitotic and asynchronous cells were resuspended in PBS + 300nM DAPI immediately after permeabilization and analyzed by FACS and microscopy. Image analysis utilized FIJI (Schindelin et al., 2012)

#### FUCCI2a FACS sorting

Prior to sorting, a partial synchronization and release was performed to enrich for cells in G1. Initially, the cells were treated as for mitotic arrest (above). After 5hrs Nocodazole treatment, cells were kept in the dish (no shake-off), gently washed 2x with PBS, and then released into pre-warmed (37°C) media for 2hrs. Cells were then collected in bulk and a BD FACS Aria II was utilized to sort cells in early G1 (mCherry− Gmn−), late G1 (mCherry+ Gmn−) and S/ G2 (Gmn+). An aliquot of each sorted population was re-checked for purity and cell cycle analysis while remaining cells were processed for pre-mRNA RT-qPCR.

#### pre-mRNA RT-qPCR

RNA was isolated using RNeasy Mini Kit (Qiagen, 74106), and DNase treated (Qiagen, 79254). cDNA was prepared using iScript Reverse Transcription Supermix (Biorad, 1708841). Real Time PCR was performed using PowerUp SYBR Green Master Mix (Applied Biosystems, A25742) and pre-mRNA primer sequences are listed in Table S5.

#### Western Blot

Cells were resuspended in 1X Laemmli Buffer. Samples were sonicated for 5 cycles using the Bioruptor Pico (Diagenode) and then boiled for 5 mins. Samples were then used for Western Blot analysis using the following antibodies: Actin HRP (ab49900), H3K27ac (ab4729), and H3K9ac (ab32129).

#### ATAC-seq

ATAC-seq was performed as previously described (Buenrostro et al., 2013). In brief, a total of 50,000 cells were washed once with 50μL of cold PBS and resuspended in 50μL lysis buffer (10 mM Tris-HCl pH 7.4, 10 mM NaCl, 3 mM MgCl2, 0.2% (v/v) IGEPAL CA-630). They were then centrifuged for 10min at 800xg at 4°C, followed by the addition of 50μL transposition reaction mix (25μL TD buffer, 2.5μL Tn5 transposase and 22.5μL ddH2O) using reagents from the Nextera DNA library Preparation Kit (Illumina #FC-121-103). Samples were then incubated at 37°C for 30min. DNA was isolated using a ZYMO Kit (D4014). ATAC-seq libraries were first subjected to 5 cycles of pre-amplification. To determine the suitable number of cycles required for the second round of PCR the library was assessed by quantitative PCR as described (Buenrostro et al., 2015) and the library was then PCR amplified for the appropriate number of cycles using Nextera primers. Samples were subject to a dual size selection (0.55×–1.5×) using SPRIselect beads (Beckman Coulter, B23317). Finally, the ATAC libraries were sequenced on an Illumina Hi-Seq (2500) platform for 50bp paired-end reads.

#### PRO-seq

PRO-seq was performed as described previously (Kwak et al., 2013), with a few adjustments. All steps were performed on ice. For each replicate per time point, 2 million cells were collected, washed briefly in PBS, and resuspended in Buffer P (10mM Tris-HCl pH 8.0, 10% glycerol, 10mM KCl, 5mM MgCl2, 250mM sucrose, 0.5mM DTT, 0.1% IGEPAL) at the cell density of 2×106 cells/ml. After 1min of incubation, 5x volume of Buffer W (10mM Tris-HCl pH 8.0, 10% glycerol, 10mM KCl, 5mM MgCl2, 250mM sucrose, 0.5mM DTT) was added to dilute the IGEPAL before centrifugation at 1000xg for 5min. Supernatant was discarded and the permeabilized cells were resuspended in Buffer F (5mM Tris-HCl pH 8.5, 40% glycerol, 5mM MgCl2, 0.5mM DTT, 5 mM DTT, 1μL/ml RNase inhibitor (SUPERaseIN, Ambion)) at a density of 2×106 cells/100μl and immediately frozen in liquid nitrogen. Samples were stored at °80°C until library preparation. PRO-seq libraries were prepared using biotin-NTP-supplemented run-ons from 0.2-0.5 × 106 permeabilized cells with Drosophila nuclei spike-in. NRO-RNA was purified using Norgen RNA isolation columns (Norgen Biotek cat. # 17200), followed by base hydrolysis on ice for 10min. After the first binding and 3’-ligation, the NRO-RNA was bound to beads and all subsequent reactions (decapping, end repair, 5’-ligation, reverse transcription) were performed as in (Mahat et al., 2016), except that the reactions were done on the beads in 20μl. All biochemical steps after the 3’-ligation were performed with rotation except for reverse transcription. After reverse transcription, cDNA was eluted twice with 25μl ddH2O by heating for 1min at 80°C, returning tubes to magnet, and collecting supernatant. Libraries were sequenced on an Illumina NEXT-seq 550 for 75bp single end reads.

#### RNA-seq

Total RNA from 300,000 asynchronous cells per replicate was prepared with TRIZOL (Life technologies #15596018) following manufacturer’s instructions. Libraries were generated by the Weill Cornell Genomics core facility using the Illumina TruSeq stranded mRNA library preparation kit (#20020594) and sequenced on an Illumina HiSeq4000 platform on SE50 mode.

#### Hi-C

In situ Hi-C was performed as described previously (Rao et al., 2014). In brief, 2 million cells (per replicate per time point) were crosslinked in 1% formaldehyde at RT for 10 minutes and quenched with 125mM glycine for 5 mins at RT. Cells were resuspended in lysis buffer (10mM Tris-HCl (pH 8.0), 10mM NaCl, 0.2% Igepal CA630 (Sigma)) and chromatin was digested overnight with 100U MboI (NEB, R0147M). Fragmented ends were labeled with biotin-14-dATP (Life Tech, 19524-016). Overnight ligation was performed using 2000U T4 DNA Ligase (NEB, M0202), and then samples were reverse cross-linked with Proteinase K (NEB, P8102). DNA was ethanol precipitated overnight, washed, and then sonicated in a Diagenode Bioruptor 300 (8 cycles, 30sec on/off, medium setting). SPRIselect beads (Beckman Coulter, B23317) were used for a 300-500 bp size selection. Aliquots were removed after digestion, ligation, sonication, and size selection and run on an agarose gel for quality control. Dynabeads MyOne Streptavidin T1 beads (Invitrogen, 65602) were used for biotin pulldown. Repair of fragmented ends was performed followed by dA-tailing. 1X NEB Quick Ligation Reaction Buffer (NEB, B2200S) was used for ligation of SeqCap Adapter Kit A (Roche, 7141530001) indexes. Library amplification was performed directly off of the T1 beads using KAPA HiFi HotStart ReadyMix and primers (KAPA, KK2620) and 8 PCR cycles. Finally, a 1X cleanup was performed using SPRIselect beads. The Hi-C libraries were sequenced on an Illumina Hi-Seq 4000 platform for 50bp paired-end reads.

#### 4C-seq

For each sample, 2 million cells were crosslinked in 1% formaldehyde at RT for 10 min and quenched with 125 mM glycine for 5 min. The cell pellets were washed twice in PBS and resuspended in 300μl lysis buffer (10mM Tris-HCl (pH 8.0), 10mM NaCl, 0.2% Igepal CA630 (Sigma I8896)), then incubated on ice for 20 min. Following centrifugation at 2500xg for 5 min at 4°C, the pellet was resuspended in 50uL of 0.5% SDS and incubated for 10 min at 65°C. SDS was quenched with 145uL ddH2O and 25uL of 10% TritonX-100 for 15 mins at 37°C. 25ul of CutSmart buffer (NEB B7204S) was added with 10ul DpnII enzyme (NEB R0543M) and the samples were incubated overnight at 37°C with 700rpm rotation. After confirming the digestion efficiency, the enzyme was inactivated by incubating at 65°C for 20 mins. The samples were then diluted with 669μl ddH2O, 120μl T4 ligation buffer (NEB B0202), 60μl 10mM ATP (NEB P0756S), 120μl 10% Triton X-100, 6μl 20mg/ml BSA and 5μl 400U/μl T4 DNA Ligase (NEB M0202) for 3h on a rotor at RT. The samples were then treated with proteinase K and reverse crosslinked overnight. Following RNAse treatment, phenol/chloroform extraction and DNA precipitation, the pellets were dissolved in 100μl of 10mM Tris pH 8. The second digestion was performed by adding 20μl 10x buffer B (Fermentas), 10μl Csp6I (Fermentas, ER0211), 80μl ddH2O, and incubating overnight at 37°C with 700rpm rotation. After confirming the second digestion efficiency, the enzyme was again inactivated. Another ligation was performed by adding 300μl T4 ligation buffer, 150μl 10mM ATP, 5μl T4 DNA Ligase, and ddH2O to 3mL and incubating overnight at 16°C. The DNA was purified by phenol/ chloroform extraction and ethanol precipitation, resulting in the 4C-template. For library preparation, primers were designed around the TSS were for each example gene according to criteria previously described (Krijger et al., 2020). Library preparation was then performed using the inverse PCR strategy described in (Krijger et al., 2020). Briefly, 200 ng of 4C-template DNA was used to PCR amplify the libraries using the Roche Expand long template PCR system (Roche, 11681842001). Primers were removed using SPRIselect beads (Beckman Coulter, B23317). A second round of PCR was performed using the initial PCR library as a template, with overlapping primers to add the P5/P7 sequencing primers and indexes. The libraries were sequenced on a HiSeq4000 in SE150 mode. All of the primer sequences can be found in Table S5.

### QUANTIFICATION AND STATISTICAL ANALYSIS

#### ATAC-seq analysis

Bowtie 2 aligner (v2.2.6) (Langmead and Salzberg, 2012) with the following parameters –very-sensitive -X 2000 was used to map raw sequencing data on mm10 genome while filtering of duplicate reads and low quality reads (Q<20) was performed with the use of picard tools (MarkDuplicates command, v2.12.2) (Picard) and samtools (v1.8) (Li et al., 2009). All reads in Blacklisted genomic regions and chrM were removed from downstream analysis. Filtered paired-end reads were corrected for tn5 insertion position at each read end by shifting +4 / −5 bp from the positive and negative strand respectively. Peak calling was performed in the merged aligned reads from all time points in order to build an accessibility genome atlas (ATAC ATLAS) for the PSC cell cycle with the use of MACS2 peak calling algorithm (v2.1.1) (Zhang et al., 2008). Visualization of sequencing data on the IGV browser (Robinson et al., 2011) was perfomed with the use of bedGraph and BigWig files generated with the use of bedtools (genomeCoverageBed) (Quinlan and Hall, 2010) and bedGraphToBigWig command. All files displayed were normalized to sequencing depth and RPM (reads per million) values were generated for each BigWig file. Identification of “retained” and “lost” accessible sites was performed with the use of intersectBed for all accessible sites in mitotic and asynchronous after peak calling with MACS2.

#### PRO-seq analysis

Sequencing of PRO-seq in single-end 50bp fragments was performed for each time point. Cutadapt (v1.4.2) (Martin, 2011) was used in order to trim adaptors from sequenced reads and keep only the first 25bp of each read. Trimmed sequencing data were aligned with the use of bowtie 2 aligner in both drosophila (Dm6 version) and mouse genome (mm10 version). All reads that mapped to ribosomal RNA were excluded from the adaptor-filtered files and the remaining reads were first mapped to the Drosophila genome and the unmapped reads were isolated and aligned to the mm10 mouse genome. All aligned files were filtered for low quality reads, duplicates, reads that were mapped in blacklisted regions and chrM. Only uniquely aligned reads on the mappable genome (Umap 24 bp) were used to calculate the percentage of aligned reads in each replicate. For downstream analysis we used all protein coding genes from ensemble GRCm38.95 version while gene expression was calculated based on the number of reads within the gene body after discarding the first and last 500bp [TSS + 500bp, TES −500bp] in order to avoid getting signal from the promoter and transcription end site which are usually rich in PRO-seq signal. Based on the number of cells used for each experiment and the mapped reads in Drosophila genome (in millions) we normalized gene expression levels after scaling all reads to the sample with the minimum sequencing depth (Table S1) and then normalized to the size of the gene (Kb). Normalized RPKM values were used for downstream analysis for more than 14,000 protein coding and long non-coding genes that were found to be expressed (normalized RPKM >−1) at least in one of the 4 time points.

#### TRE and eTRE identification

Detection of regulatory elements (TRE) was performed in the merged PRO-seq aligned reads from each timepoint with the use of dREG algorithm (Danko et al., 2015; Wang et al., 2019). All identified TREs were extended +/− 100bp from both ends and those that were included in a region +/− 1kb from any transcript were discarded from the downstream analysis. The remaining TREs were identified as enhancer TREs (eTRE) and their expression levels were calculated as in PRO-seq analysis with the only difference of the size normalization which covered the whole eTRE region.

#### RNA-seq analysis

TopHat2 (Kim et al., 2013) with default setting for paired end data was used to align the data to the mouse genome (mm10) and samtools for transforming the file formats, filtering low quality reads and sorting the paired end reads. HTseq-count (Anders et al., 2015) on mapped reads was used to calculate the raw counts per transcript and additional RPKM normalization was performed in R.

#### Hi-C analysis

HiCexplorer (v1.8) (Ramirez et al., 2018) was used to analyze the 50b paired-end reads generated for Hi-C experiments in asynchronous, mitotic cells and cells in Early and Late G1 phase. Bowtie2 aligner was used to align each pair independently and HiCexlorer tools hicBuildMatrix, hicNormalize and hicCorrectMatrix were utilized in order to construct the 20kb HiC matrices, normalized for sequencing depth and perform iterative correction method (ICE) normalization (Imakaev et al., 2012).

#### TAD analysis

Matrices in 20kb resolution were used to identify topologically associated domains (TADs) with hicFindTADs and the use of the following parameters, “--minDepth 120000 --maxDepth 420000 --step 40000 --delta 0.01--thresholdComparisons 0.01 --correctForMultipleTesting FDR”. Boundaries calculated in each replicate were merged and only boundaries with an insulation score <−0.3 were considered as TAD boundaries for that time point. TAD domain score was calculated as described in (Stadhouders et al., 2018).

#### Chromatin contact analysis

Cis-chromatin interactions were calculated for each replicate with the use of FitHiC (v1.1.0) (Ay et al., 2014) with the use of the following parameters “-r 20000 -L 40000 -U 10000000”. Only loops that were common in both replicates with a q-value < 0.01 and more than 5 contacts were scored as valid loops. Contact strength per time point was calculated based on the normalized Hi-C reads, while different kinetics of loop reestablishment was estimated after k-means clustering of the z-transformed normalized Hi-C reads across all timepoints. Enrichment analysis of previously published data with LOLA (Sheffield and Bock, 2016) was performed on the accessible sites that overlapped with the anchors of each loop cluster.

#### Virtual 4C

We used filtered Hi-C read pairs as described above and in (Di Giammartino et al., 2019) before binning and normalizing each replicate. We extracted read pairs from the 10kb bin that the read mate maps around the virtual viewpoint. We defined successive overlapping windows for each chromosome at a 10kb resolution, overlapping by 90% of their length. We then counted the second mapped read mate in all overlapping bins. Read counts were normalized to the total sequencing depth of the respective replicate. Visualization was done using normalized read counts per condition.

#### Compartment analysis

In addition to 20kb normalized matrices 100kb matrices were generated with the use of hicMergeMatrixBins in order to visualize larger chromatin structures. Compartment identification was performed with the use of Cscore algorithm (v1.1) (Zheng and Zheng, 2018) for each replicate and chromosome separately with the use of the following parameter minDis = 1000000. All compartments were assigned to active (A) and inactive (B) compartments based on gene density in each 100kb region.

#### 4C-seq analysis

All sequenced reads were demultiplexed with the use of fastx-toolkit while VP primer was trimmed with the use of fastq_trimmer. Only reads containing the RE site next to the VP were considered for analysis. Bowtie2 with very-sensitive option was used for aligning the trimmed reads (size > 40bp) to the mouse genome (mm10) while samtools was used to filter for high quality uniquely mapped reads. Smoothening of the 4C signal was performed by calculating the number of raw reads within the 10kb binned genome with a sliding window of 500bp followed by sequencing depth normalization of the cis interaction identified around the viewpoint (+/− 1MB). R and DESeq were used to depict the differential enrichment between the sliding windows and bins with an enrichment of > 3 CPM, pvalue <0.05 and fold > 2 between any two time points were scored as differential interacting regions. Significant bins per time point were merged and only interactions with more than 3 bins were scored as interaction regions. Interacting regions that were less than 1kb were discarded.

#### ChIP-seq enrichment analysis

Enrichment analysis of previously published data in ESC cells was performed with LOLA (Sheffield and Bock, 2016). For gene and loop clusters the enrichment analysis was performed on the accessible sites that overlapped with the promoter (TSS+/−2.5kb) or with either of the anchors. All the accessible sites (ATAC ATLAS) identified in the time course were used as a background set and transcription factors or histone modifications with a ‘−log10(p-value)> 3’ were considered significant and presented in the heat maps.

#### ChIP-seq analysis of published data

Retained and lost peaks for all bookmarking histone modifications and transcription factors was calculated after merging all individual replicates and corresponding inputs for GSE75066 (ESRRB), GSE122589 (OCT4, SOX2), GSE131356 (CTCF), GSE109962 (TBP), GSE92846 (KLF4, H3K27ac). All data were mapped to mm10 with bowtie2 and --local –very-sensitive-local option. Filtering of low quality mapped reads and duplicate removal was performed with samtools and picard tools while additional filtering of mm10 black-regions was done with bedtools. MACS2 with default settings was used to call peaks in both mitotic and asynchronous cells and intersection of called peaks in these 2 cell states led to the classification of retained and lost peaks for each transcription factor and histone modification.

#### Gene ontology and pathway analysis

David knowledgebase (Dennis et al., 2003) was used for annotating transcripts to biological process and signaling pathways. ENSEMBL transcript ids were used as an input. Only processes and pathways with a corrected (Benjamini) p-value <0.01 were considered significant. For assigning biological processes and pathways to genomic regions (for eTRE clusters) we used GREAT analysis software (v3.0.0) (McLean et al., 2010) with the use of the ‘Basal plus extension’ option and with 2.5kb proximal upstream, 1kb proximal downstream and distal 250kb parameters. Only processes and pathways with Hyper FDR Q-value <0.05 were considered significant.

#### Assigning genes-eTREs pairs for enrichment analysis

We paired genes and eTREs with two different approaches.

1. Distance: eTREs were assigned to genes in a distance +/− 20kb from the TSS of gene. If multiple eTREs were within +/−20kb, the closest eTRE was assigned to the gene. A total of 4779 gene-eTRE pairs were used for this analysis.
2. Chromosomal interactions: All HiChIP loops (10kb resolution) that were common in two different ESC cell lines (Di Giammartino et al., 2019) were used to pair genes and eTREs. Loops with a unique eTRE in only one of the two anchors and at least one TSS in the looped anchor were scored as eTRE-gene loops. A total of 12411 gene-eTRE pairs were used for this analysis.

## Statistical Methods

All median comparisons were performed with the use of two-sided Wilcoxon’s rank sum test. P-values and summary statistics for all variables tested are provided in Table S6. K-means with default settings (“Hartigan-Wong” algorithm) was performed in R (3.4.4 version) for Hi-C and 4C-seq using the average normalized values from the two replicate experiments for each timepoint. Two-sided Fisher’s exact test was used to calculate significance of enrichment between various groups of eTREs, genes, loops, ATAC-seq peaks, H3K27ac ChIP-seq peaks, TBP ChIP-seq peaks, and CTCF ChIP-seq peaks. In the enrichment plots, size of dots indicates significance of the enrichment while color indicates Observed over Expected ratio. Comparison of the distribution of different gene clusters around boundaries was performed in R with the use of Kolmogorov-Smirov test.

## DATA AND CODE AVAILABILITY

All genomics data (Hi-C, ATAC-seq, RNA-seq, 4C-seq, and PRO-seq) generated during this study were submitted to GEO.

---

**Figure S1.**
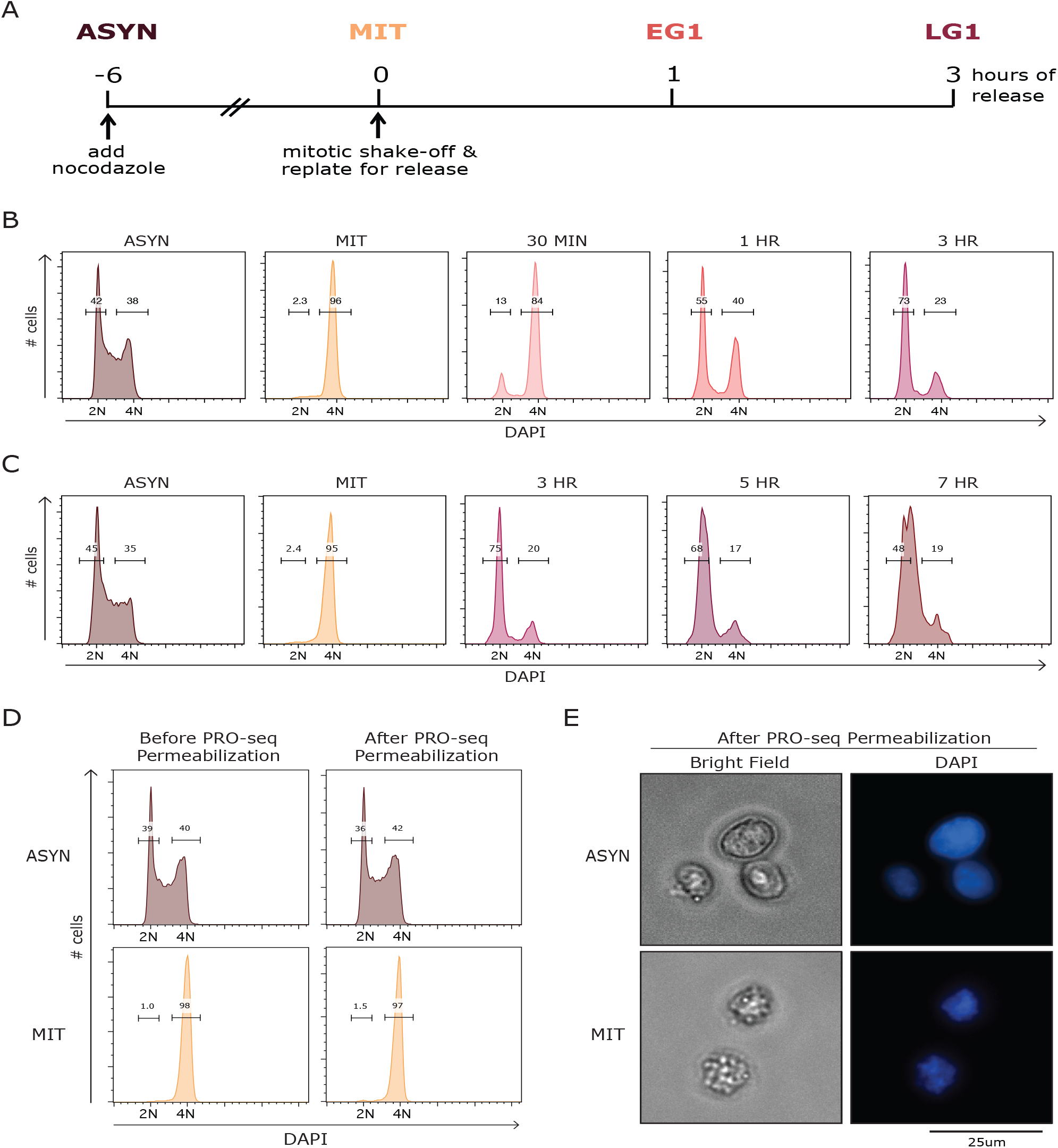
(related to Figure 1) (A) Experimental design for mitotic release time course. (B and C) Representative FACS histograms showing the percent of cells with a DNA content of 2N (G1) and 4N (G2/M) cells in mitotic release time courses with either an earlier time point (B) or later time points (C), which were used to select 1hr and 3hr as the best time points for EG1 and LG1, respectively. (D) FACS plots showing percent of cells with DNA content of 2N (G1) or 4N (G2/M) for MIT and ASYN samples both before and after PRO-seq permeabilization. (E) Representative bright field and DAPI images of MIT and ASYN cells after PRO-seq permeabilization.

**Figure S2.**
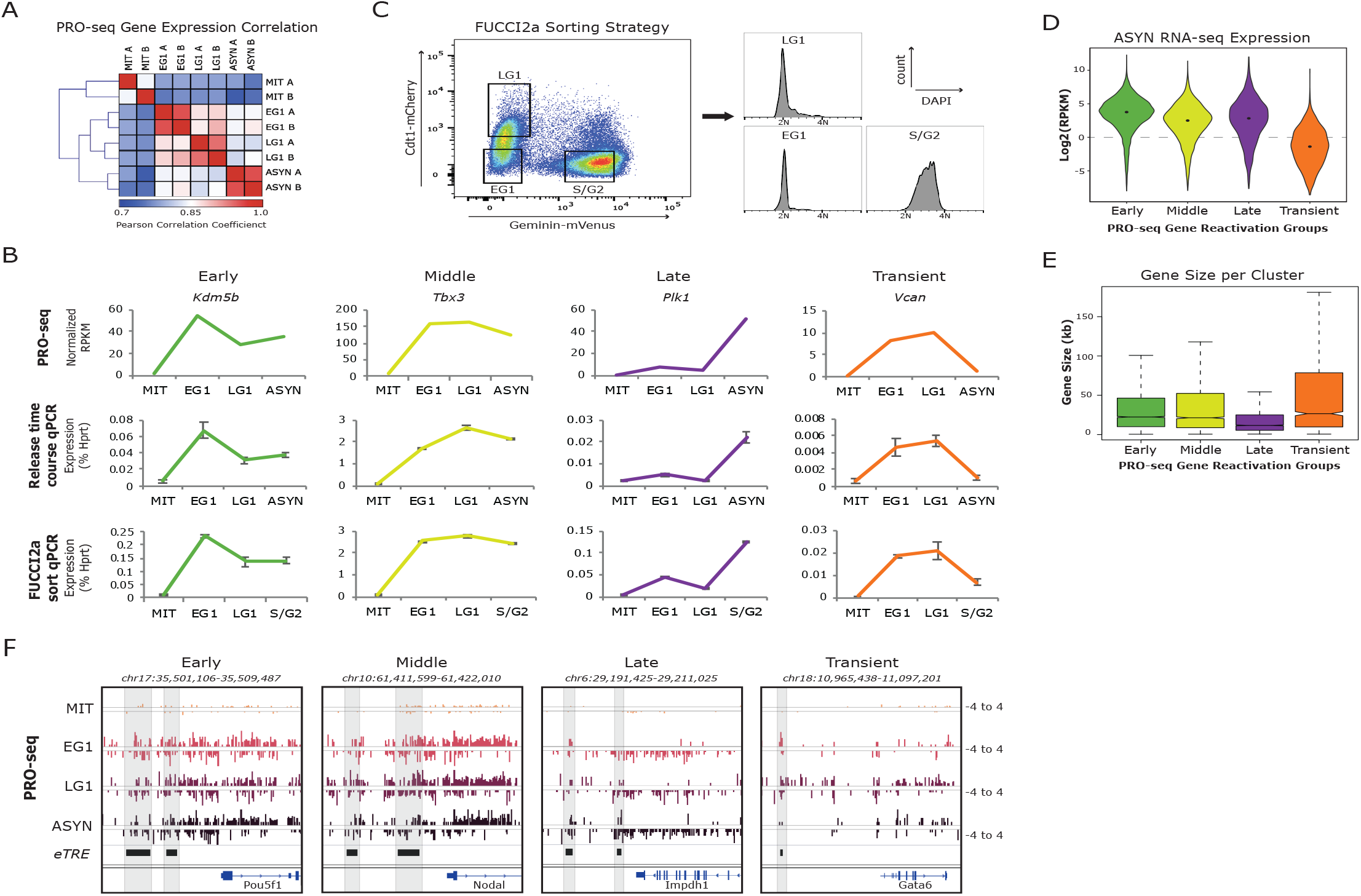
(related to Figure 2) (A) Heatmap showing the Pearson correlation among all PRO-seq samples for the 14861 protein coding genes expressed (normalized RPKM >1) in at least one time point. (B) Validation of kinetics for one example gene in each reactivation group. Top: PRO-seq expression at each time point, average of two biological replicates. Middle: pre-mRNA qPCR from two independent mitotic release time courses, error bars show +/−SEM. Bottom: pre-mRNA qPCR from two independent experiments using the FUCCI2a sorting strategy in (E), error bars show +/−SEM. MIT samples in all cases were obtained by synchronization and mitotic shake-off. (C) Sorting strategy utilizing the FUCCI2a system (Mort et al., 2014) to obtain pure populations of cells in EG1 (Cdt1−, Gmn−), LG1 (Cdt1+, Gmn−), and S/G2 (Gmn+). Prior to sorting, cells underwent a partial synchronization and release (without mitotic shake-off) to enrich for cells in G1 (see methods). Histograms show the percent of cells with a DNA content of 2N (G1) and 4N (G2/M) for each sorted population. (D)Violin plots depicting the median and density of asynchronous RNA-seq expression (Log2(RPKM)) for each gene reactivation group. See Table S6 for statistical analysis. (E) Box plots showing the median gene size (kb) for each reactivation group. See Table S6 for statistical analysis. (F) Genome browser tracks showing PRO-seq data (one replicate, plus and minus strands) for one or more eTREs from each reactivation group. Each eTRE is highlighted in gray, and the closest gene by linear distance is also shown.

**Figure S3.**
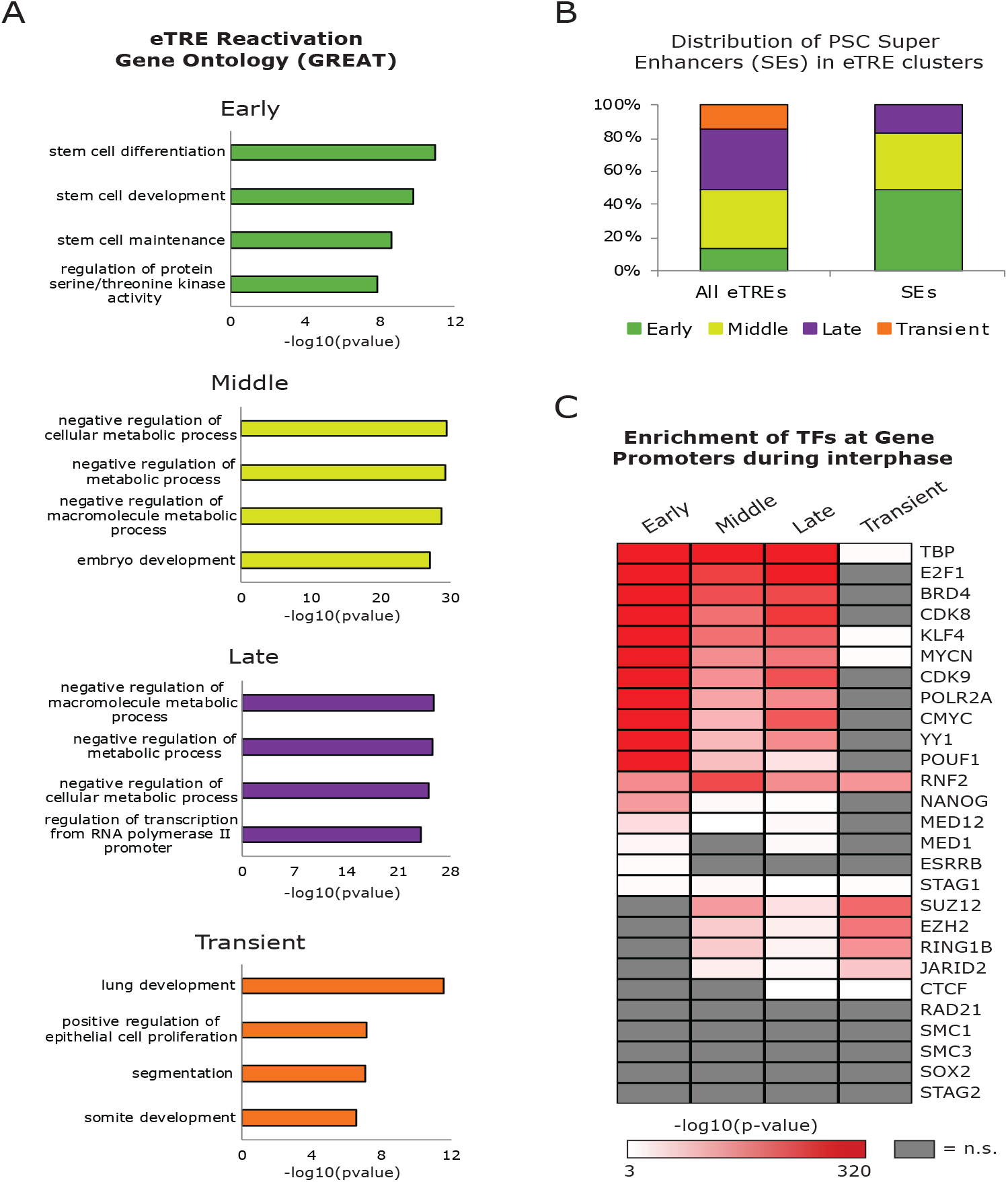
(related to Figure 3) (A) Bar plots show the enrichment (−log10(pvalue)) of the top four Gene Ontology (GO) terms for each eTRE transcriptional reactivation cluster, based on GREAT analysis (McLean et al., 2010). Corrected pvalue (Hyper FDR Qvalue) was used. (B) Distribution of PSC super enhancers (SEs), as defined by (Whyte et al., 2013), compared with all eTREs in the reactivation clusters. (C) Heat map indicates the enrichment (−log10(pvalue)) of select TFs at the accessible sites around the TSS (+/−2.5kb) of gene reactivation clusters as indicated by LOLA analysis (Sheffield and Bock, 2016). Not significant (q>0.001) terms are shown in gray.

**Figure S4.**
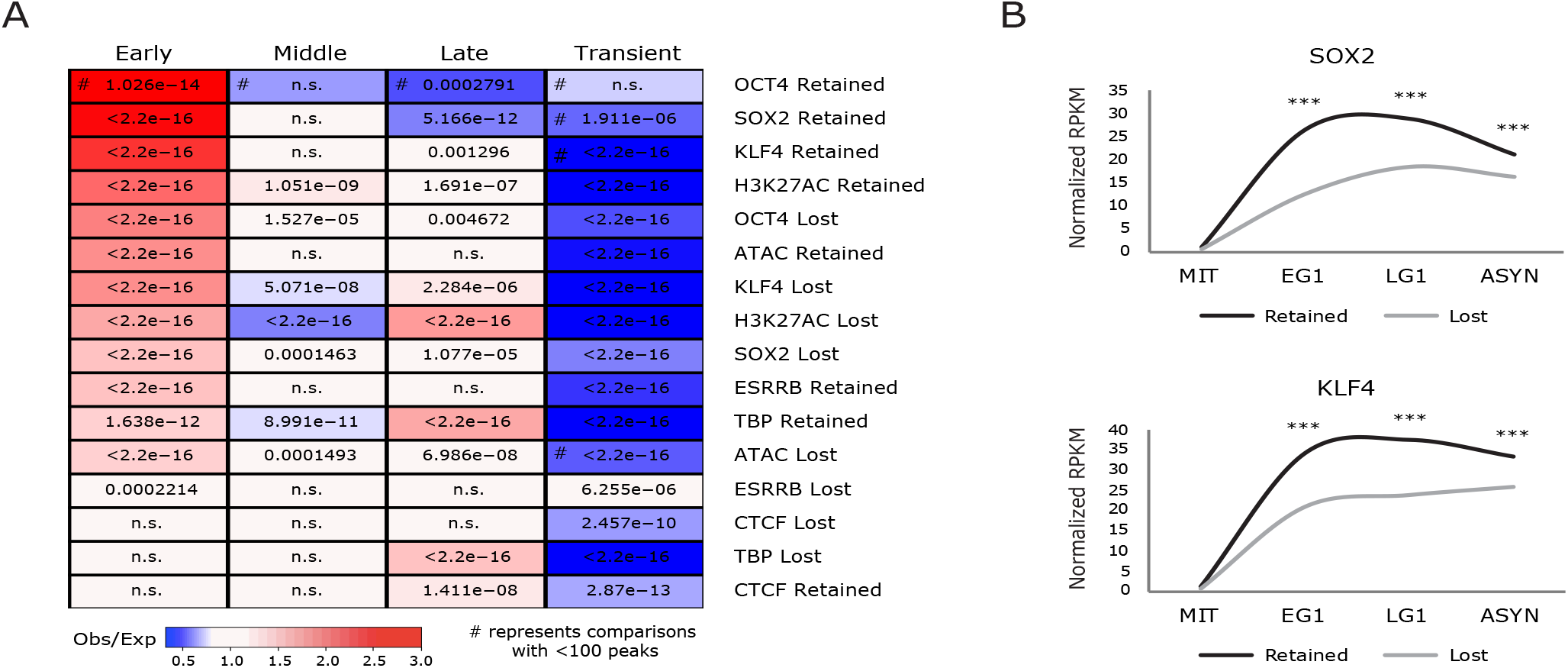
(related to Figure 4) (A) Relative enrichment or depletion of the Retained/Lost peaks for each bookmarking factor within eTRE reactivation clusters (+/−2.5kb from eTRE center, at least 1bp overlap). Color indicates ratio of observed (Obs) versus expected (Exp) frequency and p-value (two-sided Fisher’s exact test) is written in each box. Not significant (n.s.) indicates p>0.01. Comparisons using <100 overlapping peaks are denoted with a hash mark (#). See Table S3 for complete statistical analysis. (B) Median transcriptional reactivation of eTREs overlapping (at least 1bp overlap) with “Retained” and “Lost” SOX2 and KLF4 peaks. eTREs overlapping with both Retained and Lost peaks were assigned to Retained. SOX2 Retained (n=796), SOX2 Lost (n=11774). KLF4 Retained (n=1023), KLF4 Lost (n=2226). *** indicates p<0.0001, two-sided Wilcoxon’s rank sum test. See Table S6 for complete statistical analysis.

**Figure S5.**
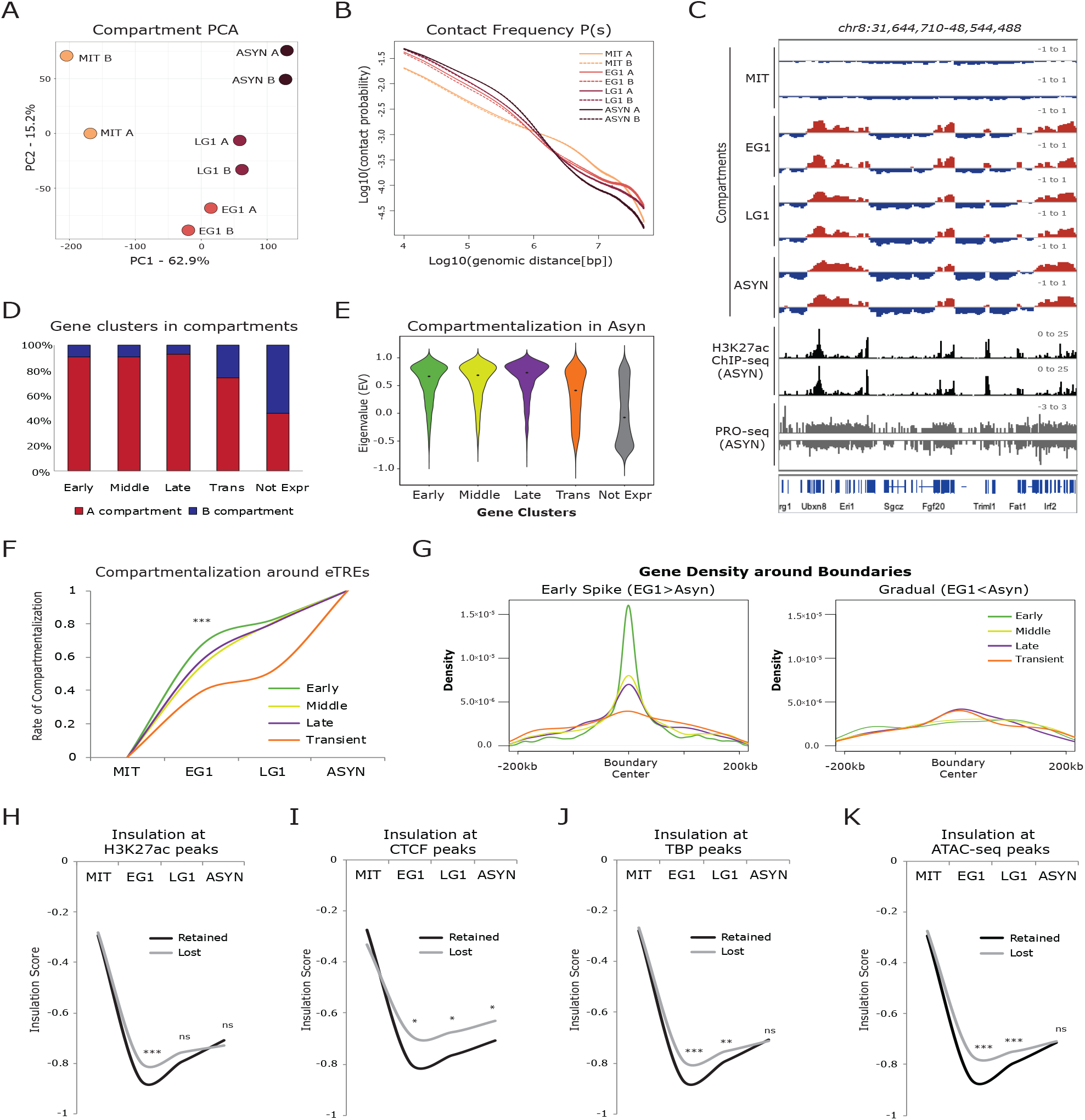
(related to Figure 5) (A) PCA of Hi-C samples based on compartment eigenvalues (EV) for each 100kb bin. (B) Contact frequency (P) versus genomic distance (s) for Hi-C datasets normalized for sequencing depth, shown on a log10 scale for both axes. (C) Genome browser tracks of a 17Mb region (chr8:31,644,710-48,544,488) showing the compartmentalization (EV) at each time point for both replicates, as well as asynchronous H3K27ac ChIP-seq (Liu et al., 2017b) and asynchronous PRO-seq tracks. (D) Bar plot showing the percent of the genes in each transcriptional reactivation cluster, as well as the 10564 not expressed (Not Expr) genes, that fall into A and B compartments. A compartments are defined as eigenvalue (EV)>0 in asynchronous cells, while B compartments are EV<0 in asynchronous cells. (E) Violin plot showing the asynchronous compartment score (eigenvalue, EV) for 100kb bins containing genes in each of the reactivation clusters (Early, Middle, Late, Transient) as well as for not expressed (Not Expr) genes. (F) Line plot showing rate of compartmentalization during mitotic exit for bins containing eTREs from the four reactivation kinetic clusters (Early, Middle, Late, Transient). If a bin contained eTREs from multiple clusters, it was prioritized as Early > Middle > Late > Transient compartments bins. *** indicates p<0.0001 for Early versus any other cluster, two-sided Wilcoxon’s rank sum test. See Table S6 for complete statistical analysis. (G) Density of the genes from each transcriptional reactivation cluster around TAD boundaries that were rapidly or gradually reset during mitotic exit. All boundaries called at any time point were pooled, and then further filtered to those that were dramatically (difference>0.2) better insulated in EG1 than ASYN (“Early Spike”, n=1772) or better insulated in ASYN than EG1 (“Gradual”, n=355). Density of the gene clusters is shown +/−200kb around the respective boundary centers. See Table S6 for statistical analysis (H-K) Median insulation score at each time point of TAD boundaries overlapping Retained or Lost bookmarking features. Only TAD boundaries called in asynchronous cells were used, and boundaries containing multiple peak types were prioritized by Retained>Lost. *s show p-value for Retained vs. Lost, two-sided Wilcoxon’s rank sum test. Not significant (ns) indicates p>0.01. (H) H3K27ac (I) CTCF (J) TBP and (K) ATAC-seq. See Table S6 for complete statistical analysis.

**Figure S6.**
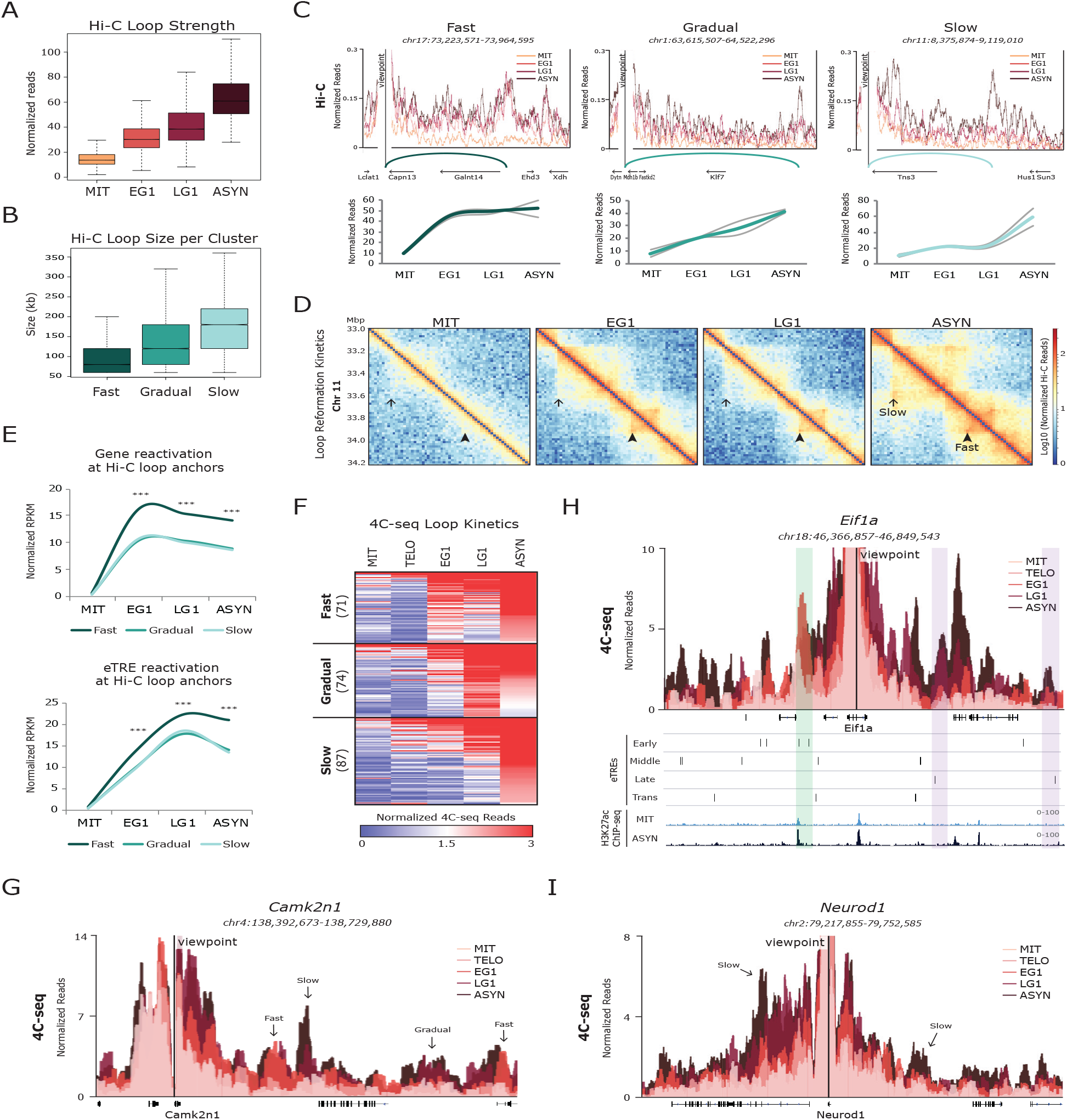
(related to Figure 6) (A) Box plot depicting the contact strength (normalized Hi-C reads) over the time course for all 52489 called loops. (B) Box plot showing median size (distance between anchors) of each Hi-C loop cluster. See Table S6 for statistical analysis. (c) Virtual 4C representations of normalized Hi-C signals around selected viewpoints to show Fast, Gradual, and Slow loop examples. Normalized Hi-C read counts of indicated loops during the time course are shown below as line plots. Individual replicates are depicted with gray lines while the average is shown in color. (D) Hi-C interaction maps (log10 Normalized Hi-C reads) of a region on chromosome 11 (chr11:33,000,000-34,200,000) at each time point. Examples of a Fast and Slow reestablished contact are indicated. (E)Line plots showing median transcriptional reactivation kinetics for genes and eTREs within Hi-C loop clusters. Anchors in multiple clusters were prioritized Fast > Gradual > Slow. Number of genes: Fast=2961, Gradual=2798, Slow=1126; and eTREs: Fast=3828, Gradual=3605, Slow=1125. *** indicates p<0.0001 for Fast versus Gradual or Slow, two-sided Wilcoxon’s rank sum test. See Table S6 for complete statistical analysis. (F) Heat map of the three K-means groups (Fast, Gradual, Slow) for all 232 loops detected by 4C-seq around 11 loci. Heat map shows average 4C-seq counts per million (CPM) over the time course for each contact. Number of loops per group is shown. (G) 4C-seq data is represented as average CPM around the viewpoint (Camk2n1 promoter), with each time point shown as an overlapping bar plot. Arrows indicate relevant contacts. (H) 4C-seq data is represented as average CPM around the viewpoint (Eif1a promoter), with each time point shown as an overlapping bar plot. Genome browser tracks underneath show the eTRE reactivation clusters (Early, Middle, Late, and Transient) and H3K27ac ChIP-seq in mitotic and asynchronous cells. A fast-reformed region with Early eTREs is highlighted in green while slower reset regions around Late eTREs are highlighted in purple. (I) 4C-seq data is represented as average CPM around the viewpoint (Neurod1 promoter), with each time point shown as an overlapping bar plot. Arrows indicate relevant contacts.

## Notes

### Competing Interest Statement

The authors have declared no competing interest.

